# A genetic link between leaf carbon isotope composition and whole-plant water use efficiency in the C_4_ grass *Setaria*

**DOI:** 10.1101/285676

**Authors:** Patrick Z. Ellsworth, Max J. Feldman, Ivan Baxter, Asaph B. Cousins

## Abstract

- Genetic selection for whole plant water use efficiency (yield per transpiration; WUE_plant_) in any crop-breeding program requires high throughput phenotyping of component traits of WUE_plant_ such as transpiration efficiency (TE_i_; CO_2_ assimilation rate per stomatal conductance). Leaf carbon stable isotope composition (δ^13^C_leaf_) has been suggested as a potential proxy for WUE_plant_ because both parameters are influenced by TE_i_. However, a genetic link between δ^13^C_leaf_ and WUE_plant_ in a C_4_ species is still not well understood.
- Therefore, a high throughput phenotyping facility was used to measure WUE_plant_ in a recombinant inbred line (RIL) population of the C_4_ grasses *Setaria viridis* and *S. italica* to determine the genetic relationship between δ^13^C_leaf_, WUE_plant_, and TE_i_ under well-watered and water-limited growth conditions.
- Three quantitative trait loci (QTL) for δ^13^C_leaf_ were found to co-localize with transpiration, biomass accumulation, and WUE_plant_. WUE_plant_ calculated for each of the three δ^13^C_leaf_ allele classes was negatively correlated with δ^13^C_leaf_ as would be predicted when TE_i_ is driving WUE_plant_.
- These results demonstrate that δ^13^C_leaf_ is genetically linked to WUE_plant_ through TE_i_ and can be used as a high throughput proxy to screen for WUE_plant_ in these C_4_ species.

## Introduction

Water availability constrains agricultural production and threatens food security in many drought-prone regions (Morison *et al.*, 2008). Therefore, improving the harvestable yield relative to water supplied to crop systems (agronomic water use efficiency; WUE_ag_) has long received attention from researchers and government agencies (Bierhuizen & Slatyer, 1965; Passioura, 1977; Sinclair *et al.*, 1984; Vadez *et al.*, 2014). It has been proposed by Passioura (1977) that yield could be improved relative to available water by increasing (1) the ratio of transpiration (*T*) to evapotranspiration (*ET*), (2) whole plant water use efficiency (ratio of biomass production to total transpiration; WUE_plant_), and (3) harvest index (HI). To date, increases in WUE_ag_ have been primarily made by improved management practices that increase T/*E*T such as reducing runoff and evaporation from the soil through improved irrigation methods (Deng *et al.*, 2006; Medrano *et al.*, 2015a), increased canopy cover (Westgate *et al.*, 1997) and mulching (Medrano *et al.*, 2015a). Additionally, selecting for greater HI has increased WUE_ag_, for example, with semi-dwarf wheat varieties (Richards *et al.*, 2014). As improvements in WUE_ag_ through management practice reach their theoretical maximum, the greatest increases in WUE_ag_ will be through improved WUE_plant_, which have so far been minimal (Condon *et al.*, 2004; Deng *et al.*, 2006; Medrano *et al.*, 2015a).

To date, the limited improvement in WUE_plant_ is primarily because WUE_plant_ is a complex trait that is influenced by 1) net CO_2_ assimilation (*A*_net_) relative to water loss via stomatal conductance (*g*_s_), (i.e. the intrinsic transpiration efficiency, *A*_net_/*g*_s_; TE_i_), 2) the proportion of carbon loss from whole plant respiration (ϕ_c_), 3) “unproductive” water loss from cuticular and nighttime transpiration (ϕ_w_), and 4) the evaporative demand between the atmosphere and the plant (See theory section; Farquhar & Richards, 1984; Farquhar, Graham D. *et al.*, 1989; Farquhar, G. D. *et al.*, 1989). Theoretically, the first three of these factors can be selected for through plant breeding, but these traits, especially ϕ_c_ and ϕ_w_, are determined by a complex set of traits that are difficult to measure and select for in breeding programs (Condon *et al.*, 2002; Flexas *et al.*, 2010; Coupel-Ledru *et al.*, 2016). Alternatively, in theory, TE_i_ is an ideal trait to select for because it is independent of environmental conditions driving changes in evaporative demands (Ghannoum, 2016), and it is an important component of WUE_plant_ because it relates to both CO_2_ and H_2_O leaf exchange, influencing both photosynthetic capacity and *T* (Condon *et al.*, 2002; Condon *et al.*, 2004). Unfortunately, the primary method of estimating TE_i_ is by gas exchange measurements of *A*_net_/*g*_s_ that do not integrate well over time and generally do not represent TE_i_ over the lifetime of the plant or even the leaf (Condon *et al.*, 2004). Furthermore, these measurements are prohibitively time-consuming and laborious, making this method impractical for selecting for WUE_plant_ in a plant-breeding program; thus, a high throughput proxy of WUE_plant_ is needed.

Alternatively, leaf carbon isotope composition (δ^13^C_leaf_) has long been promoted as a proxy for an integrated measurement of TE_i_ in C_3_ and potentially in C_4_ species (Farquhar, 1983; Farquhar & Richards, 1984; Condon *et al.*, 1987; Farquhar, Graham D. *et al.*, 1989; Farquhar, G. D. *et al.*, 1989). In C_3_ plants, the relationship between δ^13^C_leaf_ and TE_i_ has been tested and even integrated into breeding programs (Farquhar & Richards, 1984; Condon *et al.*, 2002; Condon *et al.*, 2004; Cabrera-Bosquet, L. *et al.*, 2009; Cabrera‐Bosquet *et al.*, 2011; Cabrera‐Bosquet *et al.*, 2012; Elazab *et al.*, 2012; Yousfi *et al.*, 2012; Araus *et al.*, 2013). However, it remains uncertain if δ^13^C_leaf_ is an effective proxy of TE_i_ in C_4_ species because it has not been determined if there is adequate response in δ^13^C_leaf_ to TE_i_ and if there is genotypic variation in δ^13^C_leaf_ that has a physiological relationship with WUE_plant_. Nonetheless, empirical evidence in multiple C_4_ species such as *Setaria viridis*, *S. italica*, *Zea mays*, and *Sorghum bicolor* supports the theoretical relationship between δ^13^C_leaf_ and TE_i_ (Henderson *et al.*, 1998; Cabrera‐Bosquet *et al.*, 2009; Ellsworth *et al.*, 2017). These studies also demonstrated consistent differences in δ^13^C_leaf_ between well-watered and water-limited plants that negatively correlated with TE_i_. Additionally, in *S. viridis* and *S. italica*, TE_i_ correlated with WUE_plant_ (Ellsworth *et al.*, 2017). However, a correlation between TE_i_ and WUE_plant_ has not always been found in C_3_ and C_4_ species (Terashima & Hikosaka, 1995; Gibberd *et al.*, 2001; Cerasoli *et al.*, 2004; Xu & Hsiao, 2004; Chaves *et al.*, 2007; Poni *et al.*, 2009; Tarara *et al.*, 2011; Tomás *et al.*, 2012; Tomás *et al.*, 2014; Medrano *et al.*, 2015b; Pinto *et al.*, 2015). Therefore, further investigations are needed to delineate the component traits, including TE_i_, that collectively compose WUE_plant_ and how they affect the relative importance of TE_i_ to WUE_plant_, particularly in C_4_ plants.

The genetic control of WUE_plant_ and its relationship to TE_i_ and δ^13^C_leaf_ can potentially be identified using large mapping populations grown on automated phenotyping systems that measure whole plant water use and biomass accumulation on hundreds of individual plants (Fahlgren *et al.*, 2015; Feldman *et al.*, 2018). Therefore, the physiological traits, biomass accumulation, transpiration, WUE_plant_, and δ^13^C_leaf_, can be studied to determine their genetic architecture and relationships. In fact, quantitative trait loci (QTL) have been found for δ^13^C_leaf_ in several C_3_ species such as rice (Takai *et al.*, 2006; Takai *et al.*, 2009; Xu *et al.*, 2009), barley (Teulat *et al.*, 2002), *Brachypodium distachyon* (Des Marais *et al.*, 2016), wheat (Rebetzke *et al.*, 2008), tomato (Xu *et al.*, 2008), *Arabidopsis* (Juenger *et al.*, 2005; McKay *et al.*, 2008), sunflower (Adiredjo *et al.*, 2014), soybean (Dhanapal *et al.*, 2015), cotton (Saranga *et al.*, 2004), *Quercus robur* (Brendel *et al.*, 2008), and *Stylosanthes scabra* (Thumma *et al.*, 2001). Additionally, a few studies on C_3_ plants have found co-localized QTL for δ^13^C_leaf_ and WUE_plant_ (Adiredjo *et al.*, 2014; Easlon *et al.*, 2014), and, in one case, δ^13^C_leaf_ and TE_i_ were associated with a causal gene (ERECTA; Masle *et al.*, 2005). However, to date only in two studies δ^13^C was found to be under genetic control in a C_4_ species (maize; Gresset *et al.*, 2014; Twohey III *et al.*, 2018), and in a follow-up study a genomic region was identified that affected both δ^13^C_leaf_ and WUE_plant_ (Avramova *et al.*, 2018). Now more research is necessary to develop a more thorough understanding of the physiological relationship and genetic architecture of δ^13^C_leaf_, TE_i_, and WUE_plant_, so that marker-assisted breeding can be effectively used to select for WUE_plant_ and TE_i_ in C_4_ plants.

Here a recombinant inbred line (RIL) population of 189 lines created from accession A10 of *S. viridis* (L.) P. Beauv. and accession B100 of *S. italica* (L.) P. Beauv. was used to screen for WUE_plant_, TE_i_, and δ^13^C_leaf_ (Devos *et al.*, 1998; Wang *et al.*, 1998; Feldman *et al.*, 2018). Both *S. viridis* and *S. italica* are model C_4_ grasses in the same panicoid clade as important C_4_ crops such as maize, sugarcane, sorghum, Miscanthus, and the emerging bioenergy crop switchgrass. The objectives of this study were to compare δ^13^C_leaf_ between plants grown under well-watered and water-limited conditions and to determine the genetic and physiological relationship between WUE_plant_, TE_i_, and δ^13^C_leaf_.

## Theory

Agricultural water use efficiency (WUE_ag_) can be defined as the crop yield per unit water supplied to the crop system. Where crop yield relative to water use can be calculated as a function of evapotranspiration (*ET*), the proportion of ET that is transpired (*T*/*ET*), WUE_plant_, and the harvest index (harvested proportion of biomass; Eqn 1; Passioura, 1977; Condon *et al.*, 2002) as

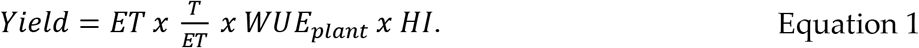

WUE_ag_ can be improved by maximizing the proportion of water inputs that are transpired at the field scale, increasing the ratio of biomass produced to water transpired, and improving the harvestable portion of the crop. Increasing the proportion of water transpired can be obtained through better crop and water management. On the other hand, the ratio of biomass and water transpired is a physiological process incapsulated in WUE_plant_ and improving WUE_plant_ is an important factor in increasing WUE_ag_ because it relates the fundamental relationship between carbon and water flux between the plant and its environment.

WUE_plant_ relates to net CO_2_ assimilation rates (*A*_net_) relative to transpiration rate (*T*) at the whole plant level, and accounts for the proportion of fixed carbon that is lost (*ϕ*_c_) such as respiration and the proportion of water loss that is “unproductive” (*ϕ*_w_) such as nighttime transpiration (*T*_night_) or cuticular evaporation because it is not associated with CO_2_ assimilation (Eqn 2; Farquhar, Graham D. *et al.*, 1989; Seibt *et al.*, 2008). The relationship of these parameters to WUE_plant_ can be defined as

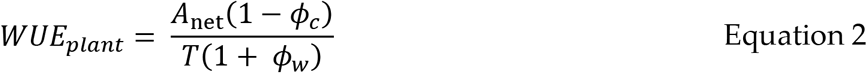

where *A*_net_ and *T* are related through stomatal conductance (*g*_s_) as

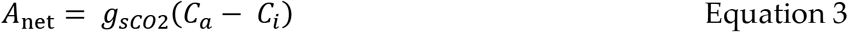

and

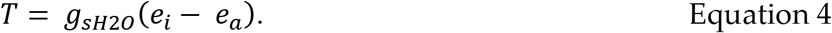

The parameter (*e*_i_ − *e*_a_) is the water vapor molar difference between intercellular and ambient air at leaf temperature, (*C*_a_ − *C*_i_) is the CO_2_ molar difference between intercellular and ambient CO_2_, and g_sCO2_ and g_sH2O_ are the conductance values for CO_2_ and H_2_O, respectively (Farquhar & Richards, 1984; Farquhar, Graham D. *et al.*, 1989; Farquhar, G. D. *et al.*, 1989). Substituting Eqns 3 and 4 into Eqn 2 gives

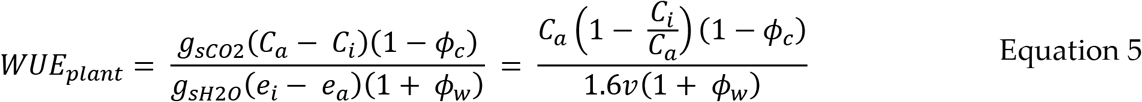

where *v* is the evaporative demand (*e*_i_−*e*_a_), and the ratio of diffusivities of H_2_O and CO_2_ in air is 1.6. The molar ratio of intercellular to ambient CO_2_ (*C*_i_/*C*_a_) influences WUE_plant_ because it represents the relative drawdown of intercellular CO_2_ (*C*_i_) by photosynthesis and the conductance of CO_2_ into the leaf and the conductance of water vapor out the leaf via the stomata. Intrinsic TE (*A*_net_/*g*_s_; TE_i_) is equal to the CO_2_ gradient from ambient to intercellular spaces (*C*_i_ − *C*_a_), which can be rewritten as *C*_a_(1−*C*_i_/*C*_a_). Therefore, Eqn 5 can be simplified as a function of TE_i_ (Eqn 6). TE_i_ and leaf carbon composition (δ^13^C_leaf_) form a relationship through their common relationship with *C*_i_/*C*_a_.

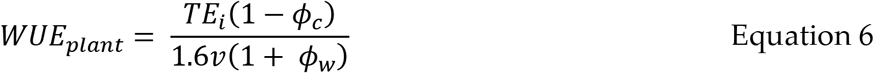

The relationship between δ^13^C_leaf_ and TE_i_ is based on 1) variation in δ^13^C_leaf_ (‰) of plants grown in the same atmospheric conditions is primarily controlled by leaf CO_2_ isotope discrimination (Δ^13^C), 2) Δ^13^C is influenced by changes in *C*_i_/*C*_a_ and 3) *C*_i_/*C*_a_, as stated above, is affected by the interrelationship *A*_net_ and *g*_s_. Therefore, TE_i_ (*A*_net_/*g*_s_) is related to *C*_i_/*C*_a_ and, in turn, Δ^13^C (Farquhar, G. D. *et al.*, 1989; Henderson *et al.*, 1998).

Finally, Δ^13^C_leaf_ is related to δ^13^C as

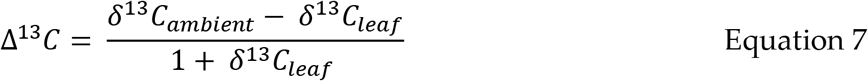

where, δ^13^C_leaf_ and δ^13^C_ambient_ are the ^13^C/^12^C ratios of the leaf and the CO_2_ in the air surrounding the leaf, respectively (Farquhar, 1983). In C_4_ species, Δ^13^C is primarily determined by fractionations associated with CO_2_ carboxylation and diffusion, the ratio of bundle sheath CO_2_ leak rate to PEP carboxylase rate (leakiness; *ϕ*), and *C*_i_/*C*_a_ (Eqn 8). Leakiness (*ϕ*) determines the slope of the relationship between Δ^13^C and *C*_i_/*C*_a_. Based on this mathematical relationship, if *ϕ* is less than 0.37, then Δ^13^C increases as *C*_i_/*C*_a_ decreases, which corresponds with increasing δ^13^C_leaf_. If *ϕ* is greater than 0.37, then the relationship reverses where Δ^13^C increases with *C*_i_/*C*_a_. In *Setaria*, *ϕ* has been found to be less than 0.37, so *C*_i_/*C*_a_ is expected to form a negative relationship with Δ^13^C and positive relationship with δ^13^C_leaf_ (Ellsworth et al. unpublished; Kubásek *et al.*, 2007). Variation in *ϕ* across genotypes could reduce or eliminate the relationship between δ^13^C_leaf_ and TE_i_ (and WUE_plant_). This is because differences in *ϕ* between plants with the same *C*_i_/*C*_a_ (and same TE_i_) would differ in carbon discrimination and δ^13^C_leaf_, while other individuals that differ in *C*_i_/*C*_a_ and *ϕ* would have the same δ^13^C_leaf_, confounding the relationship between δ^13^C_leaf_ and TE_i_. However, *ϕ* has been shown to be relatively constant across many C_4_ species and across environmental conditions such as light intensities, salinity, and CO_2_ partial pressures (Ubierna *et al.*, 2011; Sun *et al.*, 2012; Bellasio, C. & Griffiths, H., 2014; Kromdijk *et al.*, 2014; Sage, 2014; Sharwood *et al.*, 2014; Sonawane *et al.*, 2017; Sonawane *et al.*, 2018).

The relationship of Δ^13^C and *C*_i_/*C*_a_ can be defined by simplifying the relationship that was originally described by Farquhar (1984) as

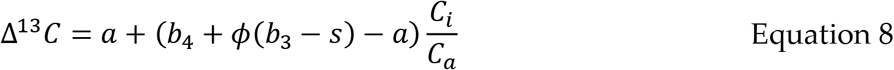

where *a* is the fractionation during diffusion of CO_2_ in air through stomata (4.4 ‰), b_4_ is the fractionations of PEP carboxylation and the preceding isotopic equilibrium during dissolution and hydration of CO_2_ (−5.7 ‰ at a leaf temperature of 25 °C) as described in (Mook *et al.*, 1974; Henderson *et al.*, 1992), b_3_ is the Rubisco fractionation (29 ‰), and *s* is the fractionation during the leakage of CO_2_ out of the bundle sheath cells (1.8 ‰) (Henderson *et al.*, 1992; Henderson *et al.*, 1998). The CO_2_ concentrating mechanism in C_4_ species reduces Rubisco isotopic discrimination against ^13^C, so that the response of Δ^13^C (and δ^13^C_leaf_) across a gradient of *C*_i_/*C*_a_ is dampened relative to that in C_3_ species (von Caemmerer *et al.*, 2014). As a result, variation in δ^13^C_leaf_ with respect to TE_i_ is related to their mutual relationship with *C*_i_/*C*_a_.

## Methods

### Plant material and growth conditions

An interspecific *Setaria* F7 recombinant inbred line (RIL) population comprised of 189 genotypes was previously generated through a cross between the wild-type green foxtail *S. viridis* accession, A10, and the domesticated *S. italica* foxtail millet accession, B100 (Devos *et al.*, 1998; Wang *et al.*, 1998; Doust *et al.*, 2009). Seeds from this population were sowed in 10 cm diameter pots pre-filled with ~470 cm^3^ of Metro-Mix 360 soil (Hummert, USA) and 0.5 g of Osmocote Classic 14-14-14 fertilizer (Everris, USA) and placed on the Bellwether Phenotyping System using an alpha lattice design replicating each genotype and treatment combination three times. Two to three replicates per genotype, including the A10 and B10 parental accessions, per treatment (1138 individuals) were transferred to the Bellwether Phenotyping System at 8 days after sowing. The experiment continued for 25 days with a photoperiod of 16 h light / 8 h night, light intensity of 500 μmol/m^2^/s, a temperature regime of 31 °C day/21 °C night and relative humidity was maintained between 40 – 80 % (Feldman *et al.*, 2018). For each replicate of each genotype, one individual plant from the well-watered treatment was grown next to a plant in the water-limited treatment. The position in the growth chamber of each paired replicates was randomly assigned and did not change during the experiment. The effect of growth chamber location was found to be negligible.

Plants were divided into two treatments: well-watered and water-limited, where soil water content was maintained at 100 % or 40 % of pot capacity (PC), respectively. Initially all seedlings were watered to pot capacity for the first two days at the Bellwether Foundation Phenotyping Facility at the Danforth Center (Feldman *et al.*, 2017; Feldman *et al.*, 2018). After two days, the potting medium in the water-limited treatment was not watered until water content dropped to 40 % PC, at which point watering resumed. To maintain soil water content at prescribed treatment levels (well-watered 100 % PC; water-limited 40 % PC), plants were watered 2 times per day until day 26 when the plants began to be watered three times per day. At this point watering took place when the light turned on, midday, and when the lights turned off. Pots were watered by weighing them and adding water until the pot weight returned to a preset weight calculated as either 100 or 40 % PC in the well-watered and water-limited treatments, respectively, as determined by Fahlgren *et al.* (2015). Prescribed soil water content across both treatment blocks was achieved by 15 days after planting. Additional detail on the experimental design and plant growth can be found in Feldman *et al.* (2017); Feldman *et al.* (2018).

### Measurements of biomass and transpiration

RGB images of individual plants were acquired using a side view camera at four different angular rotations (0°, 90° 180°, 270°) every other day, which was used to calculate biomass dry weight (Fahlgren *et al.*, 2015; Feldman *et al.*, 2018). Optical zoom was adjusted during the experiment to ensure optimal images for plant size measurements. Scaling factors relating pixel area to ground truth measurements were used to translate pixels to relative area (pixels/cm^2^). The sensitivity of the measurements limited reliable measurements of plant size and transpiration to a period from day 17 to 33 when plants were sufficiently large for accurate image analysis and measurements of transpiration (Feldman *et al.*, 2018). After the experiment concluded, 176 individual plants (91 plants from the 100 % FC and 85 from the 40 % FC) were randomly harvested and weighed for fresh aboveground biomass. Dry aboveground biomass was measured after drying at 60 °C for three days. The linear relationship between plant area from a side-view and dry aboveground biomass produced a goodness of fit similar to more complex models with multiple explanatory variables such as top-view plant area and plant height but did not overfit the data (R^2^ = 0.74). A loess function (default parameters) was used to smooth the data in the R stats library to interpolate aboveground dry biomass on an individual genotype within treatment (Chambers & Hastie, 1992). To avoid error propagation, all analyses were conducted on side-view plant area (in pixels), and after the analyses, side-view plant area was converted to dry aboveground biomass (Feldman *et al.*, 2018).

The LemnaTec instrument at the Bellwether Phenotypying Facility provided the volume of water transpired by individual plants based on gravimetric measurements and from which water use efficiency (WUE_plant_) was calculated. Total daily transpired water was the difference in total water added to the pot minus the total water added to empty pots on each calendar day. Empty pots were maintained in both the well-watered and water-limited treatments. Total water added was calculated as the difference between the measured pot weight and the weight of the pre-filled pot or the difference between current pot weight and the weight measurement on the previous day if no water was added. Empty pots were included in the experimental design to determine the volume of water lost through soil evaporation and to separate it from transpired water. Therefore, the cumulative transpiration on day 27 was the sum of all water transpired from day 17 until day 27. As described in (Feldman *et al.*, 2018), day 17 was when the difference transpired water and soil evaporation was sufficiently large to accurately measure the volume of transpired water. On days 27-33, we were able to divide plant transpiration into daytime and nighttime components because the irrigation schedule increased from two to three daily waterings, including when the lights turned on in the morning and off at night. WUE_plant_ was calculated as the ratio of dry aboveground biomass to total water transpired.

### Leaf carbon stable isotopic composition (δ^13^C_leaf_)

As a point of reference, results for genotypic and treatment effect on traits was conductance on day 27 instead of using all days and compared to δ^13^C_leaf_. The plants on day 27 were in vegetative phase and growing rapidly, typical of when physiological and gas exchange measurements would be made. Additionally, day 27 was representative in terms of QTL and other analyses conducted throughout the experiment (Feldman *et al.*, 2018). To not interrupt the experiment, the δ^13^C_leaf_ had to be collected from a harvested leaf at the end of the experiment (day 34). The leaf collected for δ^13^C_leaf_ was the youngest, uppermost, fully expanded leaf, which is the same age and development as a leaf that would have been collected on day 27 if that had been possible.

The leaves were dried at 60 °C for three days, and then 8-12 discs, having a total leaf area of 0.47 – 0.71 cm^2^, were sampled from each leaf and placed in tin capsules for stable isotopic analysis. A comparison of δ^13^C_leaf_ from sampling leaf discs versus sampling an aliquot of the completely homogenized powered leaf tissue was made on a subset of 47 leaves. The slope of δ^13^C_leaf_ from the punches regressed against δ^13^C_leaf_ from the ground leaf tissue was 0.93 ± 0.03 (R^2^ = 0.96; Fig. **S1**), and the mean difference between methods was 0.06 ± 0.04 ‰, which was similar to the IRMS precision for carbon stable isotope analysis and substantially less than the sample standard deviation of 0.5 ‰. Considering the similarity between sampling methods, all leaves were sampled using the more rapid leaf disc method.

Leaf tissue was converted to CO_2_ with an elemental analyzer (ECS 4010, Costech Analytical, Valencia, CA) and analyzed with a continuous flow isotope ratio mass spectrometer (Delta PlusXP, ThermoFinnigan, Bremen; Brenna *et al.*, 1997; Qi *et al.*, 2003). The Santrock correction was used by the IRMS software to correct for ^17^O (Santrock *et al.*, 1985). Final δ values were the mean of 5 sample peaks calibrated to the international standards NBS 19, RM 8542, and IAEA-CO-9 to calculate δ^13^C relative to Vienna Peedee belemite (V-PDB). Quality control standards were also included to determine the correction quality. Overall standard deviation for δ^13^C values was 0.07 ‰.

The stable isotope composition of carbon (δ^13^C_leaf_) was reported in δ notation,

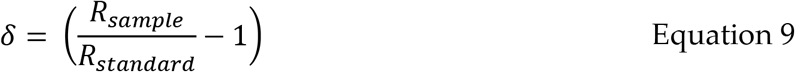

where R_sample_ and R_standard_ is the isotopic ratios of carbon (^13^C/^12^C) of the sample and international standard, respectively. The international standard used for oxygen was Vienna-PeeDee Belemite (VPDB).

### QTL analysis

The same methods used in Feldman *et al.* (2017); Feldman *et al.* (2018) were used in this study, and here the methods used are repeated. QTL mapping was performed on day 27 within each treatment group using functions in the R/qtl and funqtl packages (Kwak *et al.*, 2016). The functions were called by a set of custom Python and R scripts (https://github.com/maxjfeldman/foxy_qtl_pipeline). Two complimentary analysis methods were utilized. First, a single QTL model genome scan using Haley-Knott regression was performed to identify QTL exhibiting LOD score peaks greater than a permutation-based significance threshold (α = 0.05, n = 1000). Second, a stepwise forward/backward selection procedure was used to identify an additive, multiple QTL model based upon maximization of penalized LOD score.

The function-valued approach described by Kwak *et al.* (2016) was used to identify QTL associated with the average (SLOD) and maximum (MLOD) score at each locus throughout the experiment. Each genotypic mean trait within treatments was estimated using loess smoothing, and the QTL significance threshold was determined based upon the permutation-based likelihood of observing the empirical SLOD or MLOD test statistic. Separate, independent linkage mapping analysis performed at each time point identified a larger number of QTL locations relative to similar function-valued analysis based on the SLOD and MLOD statistics calculated at each marker. After refinement of QTL position estimates, the significance of fit for the full multiple QTL model was assessed using type III ANOVA. The contribution of individual loci was assessed using drop-one-term, type III ANOVA. The absolute and relative allelic effect sizes were determined by comparing the fit of the full model to a sub-model with one of the terms removed. QTL were grouped by finding the largest QTL by proportional variance explained and combining all QTL within 10 cM of it, then this was done for the next largest QTL and so on until all QTL were accounted for. All putative protein-coding genes (*Setaria viridis* genome version 1.1) found within a 1.5-logarithm of the odds (LOD) confidence interval were reported for each QTL in Feldman et al. (2018), which includes all QTL found in this study as well. Epistasis between QTL was evaluated using the same method as in Feldman *et al.* (2018) and Feldman *et al.* (2017) by comparing the log10 likelihood ratio of a model describing the additive effect and the additive interaction between two QTL with a model only containing the additive interaction between the two QTL. The significance threshold was determined through permutation (α = 0.05, n = 100).

### Statistical analysis

Statistical analyses were conducted in R version 3.4.0 (R Team R_Core_Team, 2013). Homogeneity was tested based on plotting predicted fit versus residuals. Using the extRemes package (version 2.0-8), normality was tested by plotting residuals on quantiles-quantiles plots. Within treatment comparisons were made on each trait using a two-factor analysis of variance (ANOVA) where the factors were treatment and genotype. Broad-sense heritability was calculated the proportion of total variance of a trait is attributed to genotypic variation 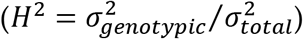. Using lmodel2 (version 1.7-2) package, model II regression analysis (standard major axis) were used instead of ordinary least squares regression for all linear regressions because neither variable was controlled, both varied naturally with their own associated error, and the physical units of both variables were not the same. Two-way ANOVAs were used to determine if the effect of treatment and genotype on each trait was significant.

## Results

### *Dry biomass, transpiration, and* WUE_plant_ *traits*

Whole plant biomass accumulation and total transpiration were analyzed throughout the experiment, and their relationship remained relatively constant through time (Fig. **S1**; Feldman *et al.*, 2018). We selected day 27 for our analysis when all genotypes were in the vegetative stage. On this day, dry biomass estimated from side view images and validated with final harvest biomass (Feldman *et al.*, 2018) varied significantly across genotypes from 0.1 to 8.33 g and 0.11 to 3.75 g in the well-watered and water-limited treatments, respectively. Cumulative transpiration ranged from 0 to 1470 ml in the well-watered and 0 to 379 ml in water-limited plants. There was a significant difference in dry biomass and transpiration rates between genotypes in both irrigation treatments and genotype x treatment effects (Fig. **1**; Table **1**), and across genotypes the water-limited treatment significantly reduced dry biomass and transpiration by 54 and 72 %, respectively (Table **1**). The ratio of dry biomass relative to the amount of total transpiration (WUE_plant_) was 58 % higher in the water-limited treatment and ranged across genotypes from 4.76 to 32.4 (g/L) in the well-watered treatment and 6.1 to 72.34 (g/L) in the water-limited treatment, respectively (Fig. **1**; Table **1**). Total transpiration was divided into daytime (*T*_day_) and nighttime (*T*_night_) components, where *T*_day_ and *T*_night_ ranged from 5 to 141 and 0 to 42 in the well-watered treatment, respectively and 2 to 69 and 0 to 30 in the water-limited treatment, respectively (Table **1**). Leaf N and C and C:N ratio were also measured and analyzed (Table **S1**and Table **S2**).

**Table 1.** Analysis of variance of traits.

| Trait | Treatment block |  |  |  |  |  | Treatment |  | Genotype |  | Treatment x Genotype |  |
| --- | --- | --- | --- | --- | --- | --- | --- | --- | --- | --- | --- | --- |
| | Well-watered | | | Water-limited | | | $F_{ddf,ndf}$ | $P$ | $F_{ddf,ndf}$ | $P$ | $F_{ddf,ndf}$ | $P$ |
| | Mean ± SE | Min | Max | Mean ± SE | Min | Max | | | | | | |
| Dry biomass (g) | 4.91 ± 0.07 | 0.10 | 8.83 | 2.21 ± 0.02 | 0.11 | 3.75 | 4968<sub>1,742</sub> | <0.001 | 15.70<sub>188,742</sub> | <0.001 | 5.71<sub>182,742</sub> | <0.001 |
| Transpiration (ml) | 565.3 ± 11.9 | 0.10 | 1470 | 159.5 ± 3.0 | 0.0 | 379.0 | 2779<sub>1,764</sub> | <0.001 | 7.468<sub>189,764</sub> | <0.001 | 3.739<sub>183,764</sub> | <0.001 |
| $T_{\text{day}}$ (ml) | 74.8 ± 1.0 | 5.0 | 141.0 | 31.9 ± 0.5 | 2.0 | 69.0 | 2692<sub>1,764</sub> | <0.001 | 5.398<sub>189,764</sub> | <0.001 | 1.745<sub>183,764</sub> | <0.001 |
| $T_{\text{night}}$ (ml) | 13.3 ± 0.3 | 0 | 42.0 | 6.8 ± 0.2 | 0 | 30.0 | 290.8<sub>1,760</sub> | <0.001 | 1.632<sub>189,760</sub> | <0.001 | 0.393<sub>183,760</sub> | 1.00 |
| WUE<sub>plant</sub> (g/L) | 9.60 ± 0.14 | 4.76 | 32.40 | 15.14 ± 0.25 | 6.06 | 72.34 | 515.8<sub>1,744</sub> | <0.001 | 6.613<sub>188,744</sub> | <0.001 | 1.063<sub>182,744</sub> | 0.29 |
| $\delta^{13}\text{C}_{\text{leaf}}$ (‰) | -13.50 ± 0.50 | -14.86 | -12.24 | -14.33 ± 0.55 | -15.99 | -12.74 | 569<sub>1,351</sub> | <0.001 | 2.054<sub>184,351</sub> | <0.001 | 1.003<sub>175,351</sub> | 0.49 |

Means ± SD of fresh biomass, transpiration, WUE<sub>plant</sub> were determined on day 27. $\delta^{13}\text{C}_{\text{leaf}}$ was collected at the end of the experiment on day 34. $T_{\text{day}}$ and $T_{\text{night}}$ are daily values for day 27, which was the first day that total transpiration could be separated into nighttime and daytime components.

**Fig. 1.**
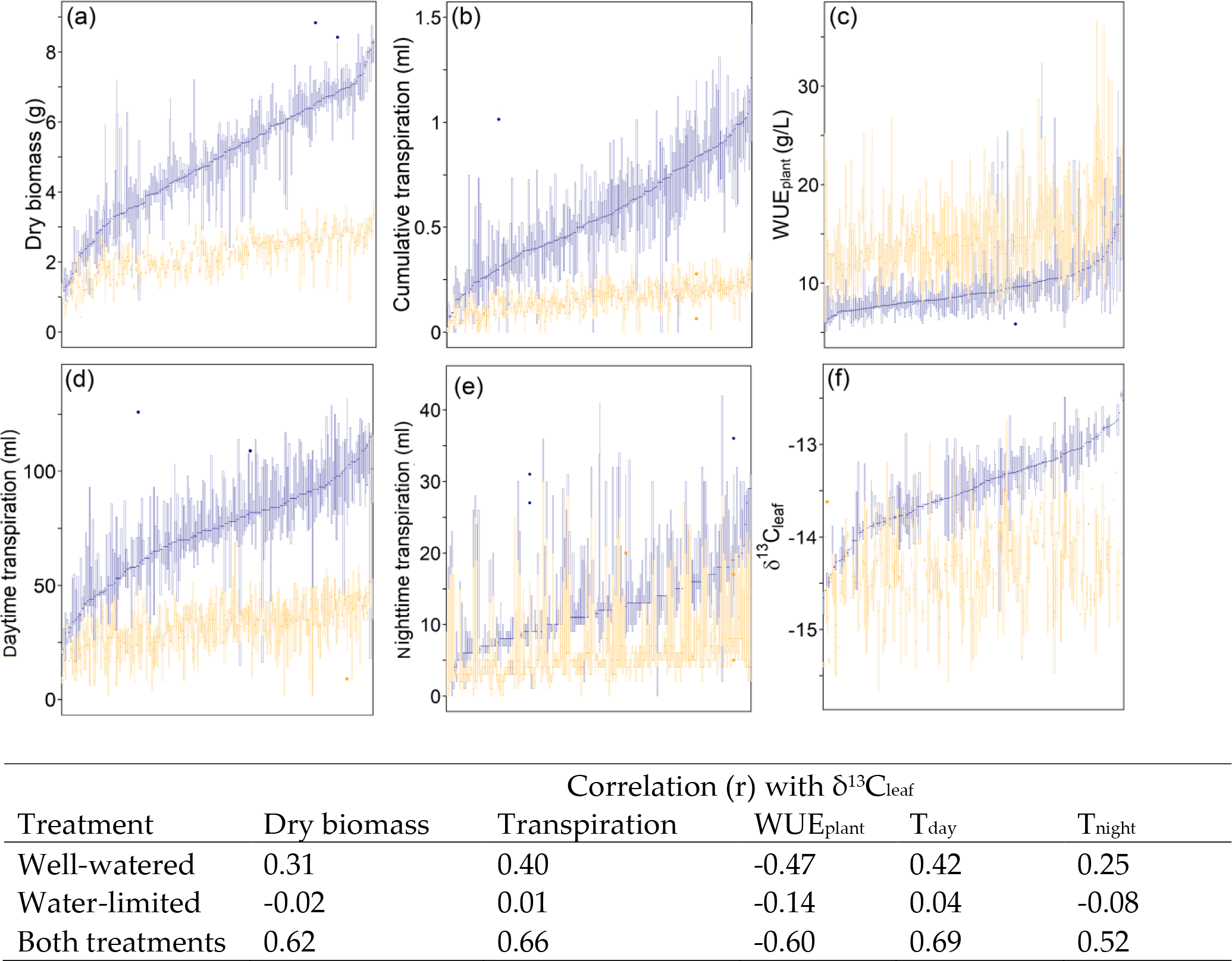
Ordered boxplots of dry biomass (A), transpiration (B), WUE_plant_ (C), *T*_day_ (D), *T*_night_ (E), and δ^13^C_leaf_ (F). All traits were measured on day 27 at peak growth, except δ^13^C_leaf_, which was measured on leaves collected at the end of the experiment on day 34. Treatment effect was significant for all traits. The table below shows the correlation coefficients of each trait from day 27 with δ^13^C_leaf_.

### *Broad-sense heritability, proportional variance and leaf carbon isotopic composition* (δ^13^C_leaf_)

In all traits, 15 - 72 % of the variance in the experiment was explained by the treatment effect (Table **2**). Additionally, in all traits, the variance ascribed to the genotype effect was relatively small but substantial given the large influence that the water limitation treatment had on these traits.

**Table 2.** Proportional variance and broad-sense heritability (H^2^) of traits on day 27 after sowing. δ^13^C_leaf_ was collected at the end of the experiment. Transpiration is cumulative transpiration throughout the experiment, while *T*_day_ and *T*_night_ are daily volumes on day 27.

| Trait | Proportional variance | | | $H^2$ | | |
| --- | --- | --- | --- | --- | --- | --- |
|  | Genotype | Treatment | G x Treatment | Both treatments | Well-watered treatment | Water-limited treatment |
| Dry biomass | 0.14 | 0.67 | 0.11 | 0.65 | 0.78 | 0.68 |
| Transpiration | 0.11 | 0.65 | 0.11 | 0.57 | 0.62 | 0.39 |
| $T_{\text{night}}$ | 0.09 | 0.72 | 0.04 | 0.43 | 0.10 | 0.00 |
| $T_{\text{day}}$ | 0.08 | 0.34 | 0.00 | 0.67 | 0.51 | 0.21 |
| $\text{WUE}_{\text{plant}}$ | 0.04 | 0.15 | 0.00 | 0.20 | 0.19 | 0.02 |
| $\delta^{13}\text{C}_{\text{leaf}}$ | 0.09 | 0.55 | 0.01 | 0.45 | 0.49 | 0.04 |

Broad-sense heritability (*H*^2^) was relatively robust for all traits, including δ^13^C_leaf_, in at least one treatment or when treatments were combined (Table **2**). For example, δ^13^C_leaf_ was significantly heritable in the well-watered treatment (0.49) but not in the water-limited treatment (0.04), which was also the case with WUE_plant_ (0.19 and 0.02 in well-watered and water-limited, respectively). In all traits, the well-watered treatment had higher *H*^2^ than the water-limited treatment. *T*_night_ had low H^2^ values for both treatments (0.1 and 0.0 in well-watered and water-limited, respectively) that limited its strength in QTL analysis but *H*^2^ was high when both treatments combined (0.43).

The δ^13^C_leaf_ values ranged from −14.7 to −12.4 ‰ in the well-watered and −15.6 to −13.2 ‰ in the water-limited treatment, with significant differences across genotypes (Fig. **2**; Table **1**). The water-limited treatment significantly reduced δ^13^C_leaf_ on average across genotypes by 0.82 ± 0.04 ‰ (Table **1**).

**Fig. 2.**
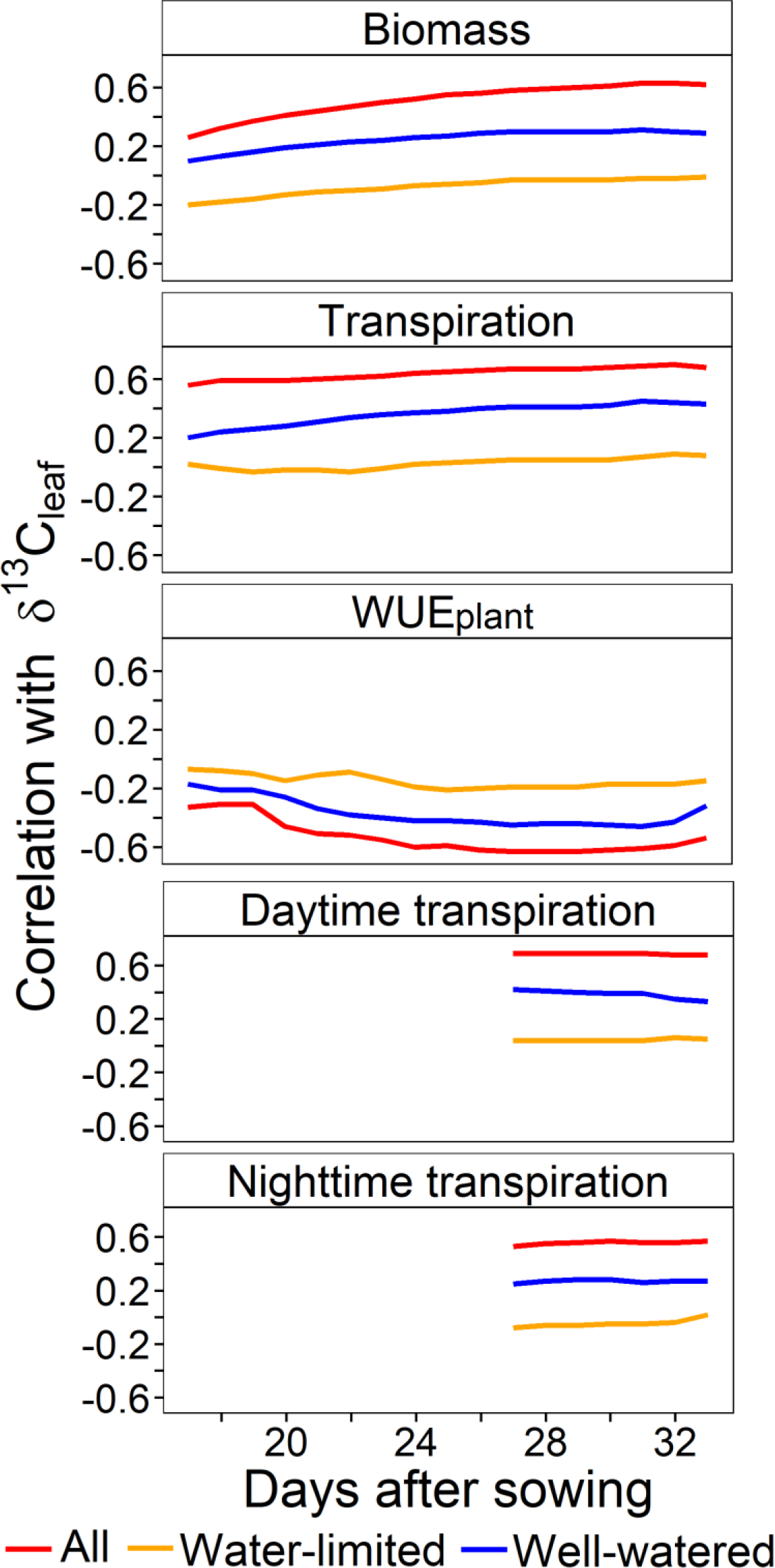
Correlation (r) of dry biomass, transpiration, WUE_plant_, *T*_day_, and *T*_night_ with δ^13^C_leaf_ through the course of the experiment. *T*_day_ and *T*_night_ were only available from day 27 to 33 because the daily irrigation schedule shifted to water when the lights turned on in the morning and when the lights turned off at night.

### Correlation of traits with δ^13^C_leaf_

Over the time course of the experiment, the correlations of δ^13^C_leaf_ with dry biomass, transpiration, WUE_plant_, *T*_day_, and *T*_night_ were relatively constant over much of the experiment, and correlations were only lower in the beginning and end of the experiment (Fig. **2**). At the midpoint of the experiment (day 27), correlations with δ^13^C_leaf_ were similar in magnitude to most days of the experiment (Fig. **S2**, **S3**, **S4**, **S5**, **S6**).

These correlations were stronger across treatments than within treatment, yet they were significant under the well-watered conditions (Fig. **2**). On day 27, the correlation coefficients of δ^13^C_leaf_ with biomass, transpiration, *T*_day_, and *T*_night_ were between 0.52 and 0.69 in magnitude across treatments and 0.25 to 0.42 for well-watered treatment (Fig. **3**). In the water-limited, correlations were low for all traits, with magnitudes ranging from −0.14 to 0.04 (Fig. **3**). On day 27, WUE_plant_ was negatively correlated with δ^13^C_leaf_ with values of −0.60, −0.47, and −0.14 for both treatments combined, well-watered, and water-limited treatments, respectively.

**Fig. 3.**
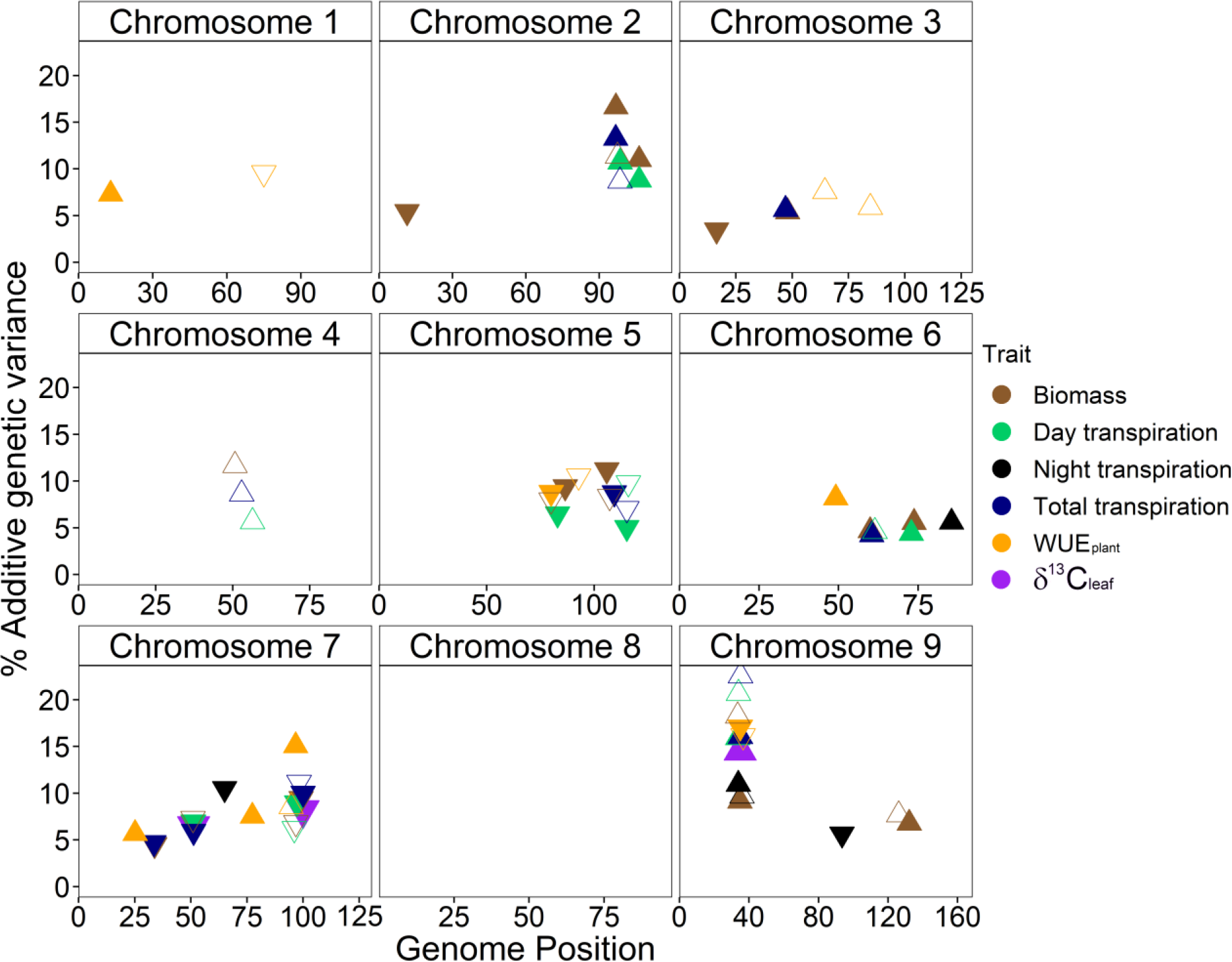
Percent additive genetic variance of QTL of whole plant traits and δ^13^C_leaf_, which was measured on leaves collected at the end of the experiment. Up-pointing and down-pointing triangles represent positive and negative mean proportional additive genetic variance, respectively. Filled and open triangles represent QTL from the well-watered and water-limited treatments, respectively.

### QTL analysis and contributions of allele composition on traits

Three QTL (chr. 7@51 centimorgans (cM), chr. 7@99 cM and chr. 9@34 cM) associated with δ^13^C_leaf_ were found in the well-watered treatment, but none were detected in the water-limited treatment (Table **3**). For simplicity, QTL were identified by their genomic position of the marker based on genetic linkage in centimorgans (cM) and was given the nomenclature of ‘chromosome @ centimorgans’. No significant epistatic interaction between QTL was detected (Table **S3**). Two of these QTL (7@99, 9@34) co-localized with WUE_plant_, dry biomass, transpiration, and *T*_day_ in both treatments on day 27. This pattern of co-localization of these two QTL is consistent over much of the experiment. QTL 7@51 was co-localized with cumulative transpiration and *T*_night_ in the well-watered treatment. Dry biomass was described in Feldman *et al.* (2018) as having this genetic architecture from days 17 through 33 (Table **3**; **S4**). Having an allele from parental accession A10 (*S. viridis*) at two of the three loci (7@51 and 7@99) increased δ^13^C_leaf_ in the well-watered treatment, while an allele from parental line B100 (*S. italica*) increased δ^13^C_leaf_ at 9@34 in both treatments (Fig. **S8**).

**Table 3.** QTL found across all traits in both treatments that are co-localized in two or more traits. Colored cell represents at least one significant QTL found in the experiment.

| Trait | Treatment | Genomic position of QTL |  |  |  |  |  |  |  |  |  |  |  |
| --- | --- | --- | --- | --- | --- | --- | --- | --- | --- | --- | --- | --- | --- |
|  |  | 2@92 | 2@113 | 3@49 | 4@50 | 5@79 | 5@104 | 6@59 | 6@75 | 7@33 | 7@51 | 7@99 | 9@34 |
| Dry biomass | well-watered | 16.6 | 11.0 | 5.4 |  | -9.4 | -11.2 | 4.7 | 5.6 | -4.5 | -6.9 | -9.4 | 9.2 |
|  | water-limited | 11.4 |  |  | 11.7 | -8.0 | -8.3 |  |  |  | -7.3 | -6.8 | 18.3 |
| Cumulative transpiration | well-watered | 13.3 |  | 5.7 |  |  | -8.7 | 4.2 |  | -4.7 | -5.9 | -10.0 | 16.1 |
|  | water-limited | 8.7 |  |  | 8.6 |  | -7.0 |  |  |  |  | -11.2 | 22.5 |
| $T_{\text{night}}$ | well-watered | | | | | | | | 5.6 | | -10.5 | | 10.9 |
|  | water-limited |  |  |  |  |  |  |  |  |  |  |  | 9.7 |
| $T_{\text{day}}$ | well-watered | 10.7 | 8.8 | | | -6.5 | -5.0 | | 4.4 | | -6.9 | -9.0 | 15.9 |
|  | water-limited |  |  |  | 5.7 |  | -9.8 | 4.6 |  |  |  | -6.2 | 20.7 |
| $\delta^{13}\text{C}$ | well-watered | | | | | | | | | | -6.5 | -8.2 | 14.5 |
|  | water-limited |  |  |  |  |  |  |  |  |  |  |  |  |
| $\text{WUE}_{\text{plant}}$ | well-watered | | | | | -8.8 | | 8.2 | | 5.7 | | 15.1 | -17.1 |
|  | water-limited |  |  |  |  | -10.5 |  |  |  |  |  | 8.6 | -16.1 |

At each marker two alleles are possible, either the allele from the parental line A10 accession of *S. viridis* (A) or the allele from the other parent, B100 accession of *S. italica* (B). The RILs in the experiment were categorized in allele classes by combining the allele (represented by letters A or B) for the three QTL associated with δ^13^C_leaf_ (in the following order: 7@51, 7@99, 9@34). The seven allele classes present in this population were AAA, AAB, ABB, BAA, BAB, BBA, and BBB, but allele class ABA was not present. For six of the seven allele classes, the SMA linear regression between dry biomass and transpiration was significant (Table **4**; Fig. **4a**). Furthermore, the regression of δ^13^C_leaf_ against the slopes of dry biomass versus transpiration showed a strong negative relationship for both the well-watered (δ^13^C_leaf_ = -0.146 slope – 12.27; R^2^ = 0.88; P = 0.006) and in the water-limited treatments (δ^13^C_leaf_ = -0.0358 slope – 13.89; R^2^ = 0.92, P = 0.0007; Fig. **4b**). Although this relationship was dampened in the water-limited treatment, it followed a similar trend (Fig. **4b**). Additionally, the order that the allele classes were positioned along the δ^13^C_leaf_ versus slope regression is very similar between treatments. In the well-watered treatment, the QTL 7@99 (represented by the second letter in three-letter allele class names) appears to have the greatest influence on this relationship, where the A10 allele was associated with a reduced slope and enriched δ^13^C_leaf_ (Fig. **4b**). Alternatively, in the water-limited treatment, the effect of QTL 7@99 on this relationship was reduced relative to the well-watered treatment. Additionally, in both treatements the mean dry biomass and transpiration for each of these allele classes had a strong significant positive relationship with δ^13^C_leaf_ (Fig. **5a** and Fig. **5b**). Similar to the transpiration versus δ^13^C_leaf_, *T*_day_ and *T*_night_ formed significant relationships with δ^13^C_leaf_, but *T*_night_ formed a weaker relationship with δ^13^C_leaf_ (Fig. **6a** and Fig. **6b**).

**Table 4.** δ^13^C_leaf_ and the regression slope of the relationship between biomass and transpiration at the allele class level.

| Allele class | Well-watered treatment |  |  |  |  | Water-limited treatment |  |  |  |  |
| --- | --- | --- | --- | --- | --- | --- | --- | --- | --- | --- |
| | Slope | R<sup>2</sup> | P | n | $\delta^{13}\text{C}_{\text{leaf}}$ | Slope | R<sup>2</sup> | P | n | $\delta^{13}\text{C}_{\text{leaf}}$ |
| AAA | 7.02 ± 0.63 | 0.88 | < 0.0001 | 26 | -13.7 ± 0.08 | 13.01 ± 1.55 | 0.77 | < 0.0001 | 26 | -14.36 ± 0.07 |
| AAB | 7.11 ± 0.46 | 0.78 | < 0.0001 | 64 | -13.15 ± 0.04 | 11.08 ± 0.87 | 0.62 | < 0.0001 | 64 | -14.28 ± 0.05 |
| ABA | Not present in RIL population |  |  |  |  |  |  |  |  |  |
| ABB | 7.71 ± 3.04 | 0.53 | 0.16 | 7 | -13.8 ± 0.07 | 11.79 ± 1.63 | 0.93 | 0.008 | 7 | -14.24 ± 0.16 |
| BBB | 12.09 ± 1.33 | 0.90 | < 0.0001 | 13 | -13.91 ± 0.07 | 14.00 ± 2.68 | 0.74 | 0.003 | 13 | -14.42 ± .13 |
| BBA | 11.71 ± 2.44 | 0.78 | 0.008 | 10 | -14.0 ± 0.09 | 20.70 ± 3.24 | 0.90 | 0.004 | 10 | -14.60 ± 0.10 |
| BAB | 7.14 ± 0.30 | 0.90 | < 0.0001 | 68 | -13.44 ± 0.04 | 10.97 ± 0.73 | 0.74 | < 0.0001 | 68 | -14.23 ± 0.05 |
| BAA | 9.50 ± 0.85 | 0.84 | < 0.0001 | 28 | -13.79 ± 0.09 | 45.56 ± 7.96 | 0.46 | 0.0005 | 28 | -14.43 ± 0.08 |

The allele class 'ABA' was not present in this RIL population. These slopes ± SEM are from the relationship found in figure 4a and are also plotted against $\delta^{13}\text{C}_{\text{leaf}} \pm \text{SEM}$ in figure 4b.

**Fig. 4.**
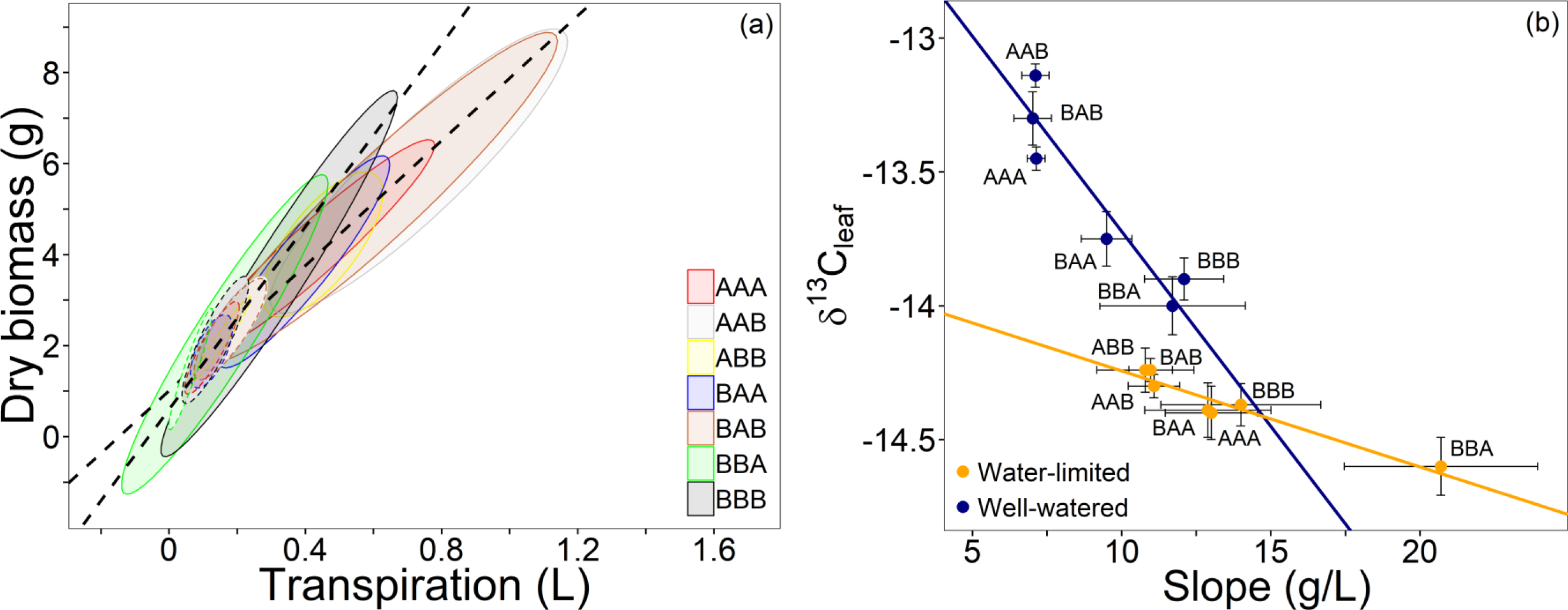
The effect of allele class on dry biomass, transpiration, and δ^13^C_leaf_. In panel (**a**), QTL 7@51, 7@99, and 9@34 were combined to produce seven alelle classes where the first letter represents the allele at QTL 7@51, the second letter represents the allele at QTL 7@99, and the third letter represents the allele at QTL 9@34. The letter ‘A’ represents the allele from the A10 parental accession (*Setaria viridis*), and ‘B’ represents the allele from the B100 parental accession (*Setaria italica*). Ellipses represent 95 % confidence intervals for the relationship of dry biomass and transpiration, and the slope of this relationship for each allele class was significant, except for allele class ‘ABB’ in the well-watered treatment (P < 0.0001). In panel (**b**), δ^13^C_leaf_ ± SEM is regressed against the slope of relationship ± SEM in panel (**a**), excluding the non-significant slope for ‘ABB’. The slope is the WUE_plant_ for an entire allele class. The regression for δ^13^C_leaf_ versus slope was significant in the well-watered (δ^13^C_leaf_ = −0.146 slope – 12.27; R^2^ = 0.88; P = 0.006) and in the water-limited treatments (δ^13^C_leaf_ = -0.0358 slope – 13.89; R^2^ = 0.92, P = 0.0007).

**Fig. 5.**
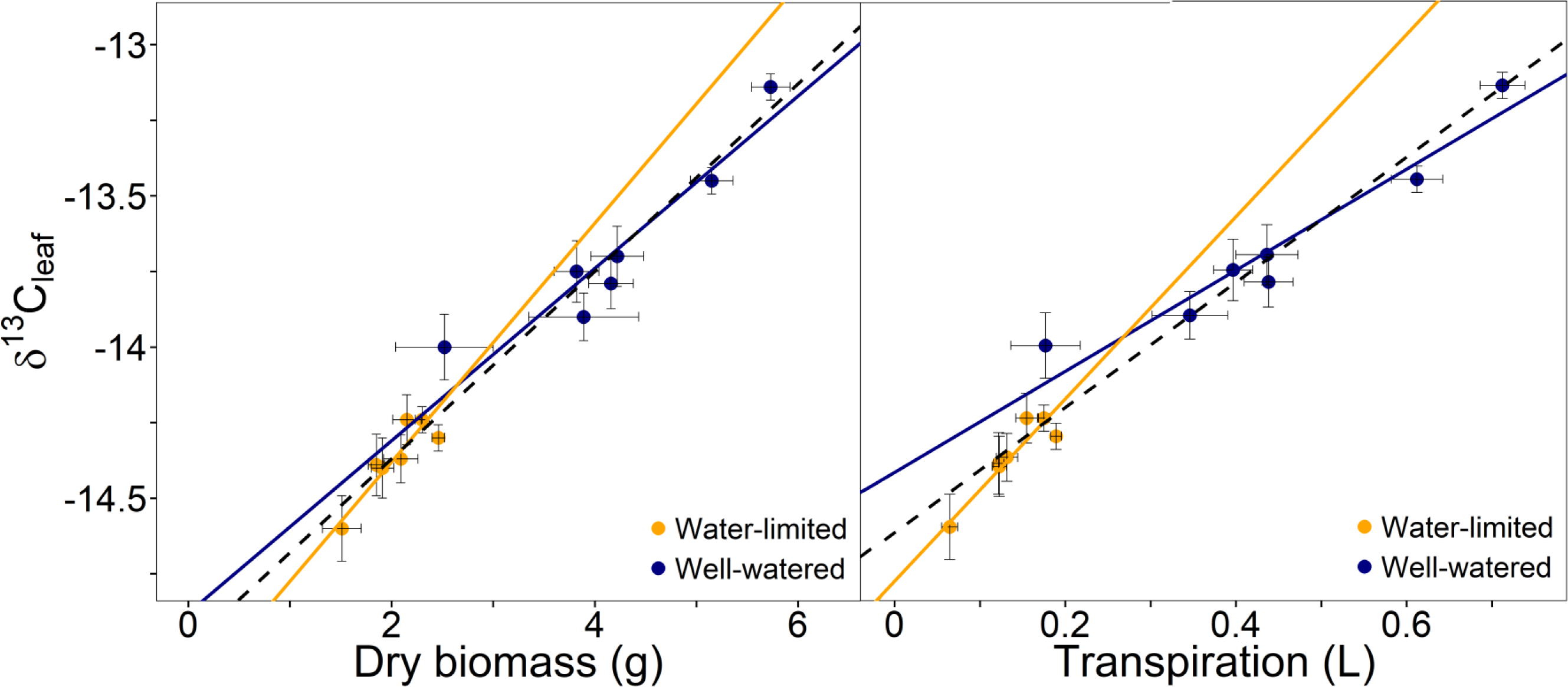
The effect of allele class on the relationship of dry biomass and cumulative transpiration with δ^13^C_leaf_. Like in Fig. 4, QTL 7@51, 7@99 and 9@34 were combined to produce seven alelle classes. In panel **a**, the mean δ^13^C_leaf_ was regressed against dry biomass for each treatment and both combined (δ^13^C_leaf_ = 0.285 dry biomass – 14.88; R^2^ = 0.87; P = 0.002 for well-watered; δ^13^C_leaf_ = 0.395 dry biomass – 15.17; R^2^ = 0.78; P = 0.008 for water-limited; δ^13^C_leaf_ = 0.310 dry biomass – 14.99; R^2^ = 0.96; P < 0.0001 for both treatments combined). In panel **b**, δ^13^C_leaf_ is regressed against cumulative transpiration for treatment and both treatments combined (δ^13^C_leaf_ = 1.67 transpiration – 14.42; R^2^ = 0.92; P = 0.0006 for well-watered; δ^13^C_leaf_ = 3.016 transpiration – 14.78; R^2^ = 0.85; P = 0.003 for water-limited; δ^13^C_leaf_ = 2.07 transpiration – 14.62; R^2^ = 0.95; P < 0.0001 for both treatments). Regression of both treatments combined is identified by black, dashed line.

**Fig. 6.**
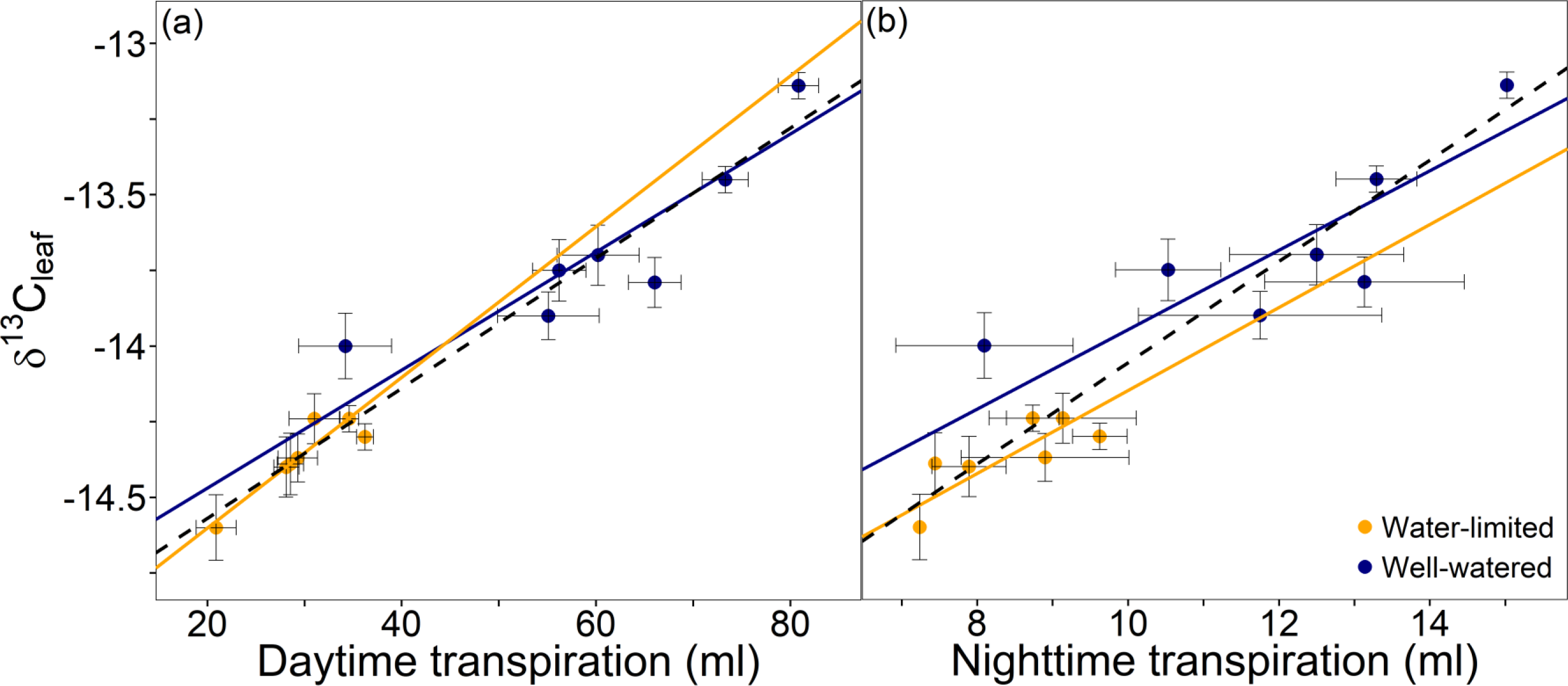
The effect of allele class on the relationship of *T*_day_ and *T*_night_ with δ^13^C_leaf_. Like in Fig. 4, QTL 7@51, 7@99 and 9@34 were combined to produce seven alelle classes. In panel (**a**), the mean δ^13^C_leaf_ was regressed against *T*_day_ and *T*_night_ for each treatment and both combined (δ^13^C_leaf_ = 0.0195 *T*_day_ – 14.86; R^2^ = 0.78; P = 0.008 for well-watered; δ^13^C_leaf_ = 0.0249 *T*_day_ – 15.10; R^2^ = 0.81; P = 0.006 for water-limited; δ^13^C_leaf_ = 0.0215 *T*_day_ – 15.00; R^2^ = 0.93; P < 0.0001). In panel (**b**), δ^13^C_leaf_ is regressed against *T*_night_ for each treatment and both treatments combined (δ^13^C_leaf_ = 0.131 *T*_night_ – 15.26; R^2^ = 0.67; P = 0.02 for well-watered; δ^13^C_leaf_ = 0.137 *T*_night_ – 15.52; R^2^ = 0.60; P = 0.04 for water-limited; δ^13^C_leaf_ = 0.167 *T*_night_ – 15.73; R^2^ = 0.85; P < 0.0001 for both treatments). Regression of both treatments combined is identified by black, dashed line. These values of transpiration represent transpiration on day 27 only because day 27 was the first day that T_day_ and T_night_ could be separated.

## Discussion

Leaf carbon isotope composition (δ^13^C_leaf_) has been theoretically related to TE_i_ (*A*_net_/*g*_s_) for both C_3_ and C_4_ species (Farquhar & Richards, 1984; Condon *et al.*, 1987; Henderson *et al.*, 1998; Condon *et al.*, 2002; Condon *et al.*, 2004). Despite the potential dampening effect that the CO_2_-concentrating mechanism has on δ^13^C_leaf_ variability in C_4_ plants (von Caemmerer *et al.*, 2014), δ^13^C_leaf_ in the *Setaria* RIL population presented here exhibited a significant genetic (range of 2.4 ‰) and environmental (mean difference of 0.82 ± 0.04 ‰ between treatments) effect. These results also show considerable genotype by treatment response, consistent with previous studies of well-watered and water-limited C_4_ plants (Monneveux *et al.*, 2007; Cabrera-Bosquet, Llorenç *et al.*, 2009; Ellsworth *et al.*, 2017). In the well-watered treatment, the δ^13^C_leaf_ had relatively strong correlations with WUE_plant_ and its component traits, biomass and transpiration. These correlations were consistent throughout much of the experiment and corroborate the theoretical relationship between δ^13^C_leaf_ and WUE_plant_ through TE_i_. However, in the water-limited plants the lack of significant correlations between δ^13^C_leaf_ and other traits (e.g. WUE_plant_, biomass, and transpiration) was likely due to restricted stomatal conductance (*g*_s_) across most genotypes, minimizing individual differences in TE_i_, similar to what was found in C_3_ species (Lambrides *et al.*, 2004; Adiredjo *et al.*, 2014).

These data demonstrate a significant genetic and environmental influence on δ^13^C_leaf_ in a C_4_ species related to differences in WUE_plant_. These relationships are further supported by the fact that δ^13^C_leaf_ shared a similar genetic architecture with WUE_plant_ and its component traits. For example, QTL (7@51, 7@99, 9@33) found for δ^13^C_leaf_ are pleiotropic loci, co-localized with leaf composition traits and whole plant traits such as biomass production, transpiration, and WUE_plant_. In the current study, WUE_plant_ was calculated differently from Feldman *et al.* (2018); however, the principal QTL for WUE_plant_ and its component traits remained similar between studies. The one notable difference was that the most significant QTL for WUE_plant_ calculated with transpiration (T) instead of ET shifted to 7@99 and 9@33. This highlights the apparent importance of using T in linking δ^13^C_leaf_ to WUE_plant_. This rationale is supported by theories describing the relationship between WUE_plant_ and δ^13^C_leaf_, suggesting that in this RIL population TE_i_ has a greater influence on WUE_plant_ than the other components *ϕ*_w_, *ϕ*_c_, and *r* described in Theory section.

An additional trait that might influence the relationship between δ^13^C_leaf_ and WUE_plant_ is bundle sheath leakiness (*ϕ*), where *ϕ* is defined as the ratio of bundle sheath CO_2_ leak rate to the rate of PEP carboxylase. Changes in *ϕ* influence the relationship between Δ^13^C_leaf_ and CO_2_ availability (*C*_i_/*C*_a_) such that differences in δ^13^C_leaf_ can result from variation in *ϕ* instead of TE_i_. However, in this experiment it is unlikely that *ϕ* is the primary driver of δ^13^C_leaf_. First, *ϕ* would have to be heritable to explain the genotypic effect in δ^13^C_leaf_. Although this is possible there are no studies, that we are aware of, showing *ϕ* as heritable and under genetic control. Second, the consistent depletion in δ^13^C_leaf_ in response to water limitation could occur if *ϕ* increased across all genotypes in the water-limited treatment. However, several studies have failed to find significant differences in *ϕ* under various environmental growth conditions including light gradients, salinity, and water limitation across species (Ubierna *et al.*, 2011; Bellasio, Chandra & Griffiths, Howard, 2014a; Bellasio, Chandra & Griffiths, Howard, 2014b; Bellasio, C. & Griffiths, H., 2014; Sharwood *et al.*, 2014; Sonawane *et al.*, 2017; Sonawane *et al.*, 2018; Sonawane and Cousins unpublished results). Third, *ϕ*-driven variation in δ^13^C_leaf_ cannot explain the strong negative correlation between δ^13^C_leaf_ and WUE_plant_ because increasing *ϕ* decreases the efficiency of the carbon-concentrating mechanism through overcycling (von Caemmerer *et al.*, 2014). This would decrease the photosynthetic efficiency, which, in turn, decreases TE_i_ and WUE_plant_. Finally, the similar genetic architecture between δ^13^C_leaf_ and WUE_plant_, biomass and transpiration would not be expected if the variation in δ^13^C_leaf_ was driven primarily by *ϕ* but rather TE_i_. Therefore, variation in *ϕ* across individual plants could certainly contribute to δ^13^C_leaf_, potentially reducing the strength of the relationship between δ^13^C_leaf_ and WUE_plant_. Nonetheless, in this experiment there was a strong relationship between δ^13^C_leaf_ and WUE_plant_ and its component traits.

Theoretical and empirical experiments indicate that δ^13^C_leaf_ and TE_i_ should be negatively correlated (Farquhar, 1983; Henderson *et al.*, 1998; von Caemmerer *et al.*, 2014; Ellsworth *et al.*, 2017). Therefore, if TE_i_ is a strong component of WUE_plant_, then there should also be a negative relationship between δ^13^C_leaf_ and WUE_plant_, as seen in the data presented here. In this RIL population there was a stronger relationship of δ^13^C_leaf_ and WUE_plant_ across allele classes based on the three QTL for δ^13^C_leaf_. The strong negative relationship between δ^13^C_leaf_ and WUE_plant_ across these allele classes further supports the link of TE_i_ between δ^13^C_leaf_ and WUE_plant_. Across these allele classes, the relationship of δ^13^C_leaf_ and WUE_plant_ is also seen in the water-limited treatment even though there were no QTL detected for δ^13^C_leaf_ under this treatment. This is likely due to the dampened response of δ^13^C_leaf_ to water-limited conditions as seen with previous studies (Avramova *et al.*, 2018). However, given that the water limitation did not remove the underlying relationship between TE_i_ and WUE_plant_, the inability to detect QTL is likely due to the reduced variation in δ^13^C_leaf_, which reduced the magnitude of the genotypic response and decreased the signal to noise ratio. Nonetheless, the basic relationship between δ^13^C_leaf_ and WUE_plant_ remained in the water-limited treatment, as did the relative order of the allele classes. For example, under both well-watered and water-limited conditions the allele classes AAB and BAB had the lowest WUE_plant_ and most enriched δ^13^C_leaf_.

Whereas allele classes AAA and BAA were in the middle in both traits, and allele classes BBB and BBA had the highest WUE_plant_ and most depleted δ^13^C_leaf_ under both treatments. This trend indicates a strong allelic effect on relationship between δ^13^C_leaf_, TE_i_, and WUE_plant_ that may allow δ^13^C_leaf_ to be used as a proxy for TE_i_, WUE_plant_ in well-watered and water-limited conditions.

### Conclusion

WUE_plant_ is driven by a balance between carbon assimilation and water lost via stomates (*TE*_i_) and other whole plant processes such as *ϕ*_c_, *ϕ*_w_, and *r* (when only aboveground biomass is measured). Hence, the relationship between δ^13^C_leaf_ and WUE_plant_ is only apparent if TE_i_ has a strong influence on WUE_plant_. In this C_4_ grass RIL population, δ^13^C_leaf_ showed a significant and consistent response to water limitation, significant genotypic variation, and significant heritability. Additionally, δ^13^C_leaf_ correlated with transpiration, biomass, and WUE_plant_, suggesting a physiological relationship among these traits. This is further supported by the fact that there were even stronger negative correlations between δ^13^C_leaf_ and WUE_plant_ within the allele classes defined by QTL of δ^13^C_leaf_. This suggests that differences in TE_i_ is driving the differences in both WUE_plant_ and δ^13^C_leaf_. This relationship between δ^13^C_leaf_ and WUE_plant_ across allele classes emphasizes the intrinsic role of TE_i_ in this relationship and implies that δ^13^C_leaf_ can detect variation in WUE_plant_ when TE_i_ is a major driver of WUE_plant_. The outcome of this research demonstrates that δ^13^C_leaf_ has the potential to screen for TE_i_ in a marker-assisted C_4_ plant breeding program. However, additional work is needed to better understand the genetic controls of δ^13^C_leaf_, TE_i_ and WUE_plant_. Furthermore, research is needed to explore the use of δ^13^C_leaf_ in other C_4_ species and under field settings to better understand the complex interaction of traits and causal genes that influence WUE_plant_, TE_i_, and δ^13^C_leaf_.

## Acknowledgements

This project was funded by U.S. Department of Energy (award number DE-SC0008769) to ABC and IB. ABC is supported in part by Meyer Distinguished Professorship. IB was supported by the US Department of Agriculture—Agricultural Research Service.

## Author contributions

M.F. and I.B. performed the experiment; P.E and M.F. analyzed the results; P.E. wrote the manuscript; M.F., A.C., and I.B. contributed to the development and writing of the manuscript.

## Appendix I Glossary of terms

| Term | Definition |
| --- | --- |
| $\Delta^{13}\text{C}$ | Photosynthetic carbon discrimination ($\delta^{13}\text{C}_{\text{ambient}} - \delta^{13}\text{C}_{\text{leaf}}$) |
| $\delta^{13}\text{C}_{\text{leaf}}$ | Leaf carbon isotopic composition (‰) |
| $\delta^{13}\text{C}_{\text{ambient}}$ | Isotopic composition of ambient $\text{CO}_2$ (‰) |
| $\text{WUE}_{\text{ag}}$ | Agricultural water use efficiency (crop yield/water applied to crop) |
| $\text{WUE}_{\text{plant}}$ | Ratio of biomass accumulated per water transpired |
| $\text{TE}_i$ | Intrinsic transpiration efficiency ($A_{\text{net}}/g_s$) |
| HI | Harvestable index |
| $C_i/C_a$ | Intercellular to ambient $\text{CO}_2$ concentration |
| $g_s$ | Stomatal conductance |
| $A_{\text{net}}$ | Net photosynthetic rate |
| $T$ | Transpiration rate |
| ET | Evapotranspiration |
| $V$ | Evaporative demand; ($e_i - e_a$) |
| $e_i - e_a$ | Water vapor molar difference between intercellular and ambient air at leaf temperature |
| $\phi_w$ | Proportion of water used by plant that is unproductive water loss (e.g. nighttime and cuticular transpiration) |
| $\phi_c$ | Proportion of fixed carbon lost through respiration |
| $R$ | Proportion of biomass |
| $A$ | Fractionation during diffusion of $\text{CO}_2$ in air through stomata (4.4 ‰) |
| $T_{\text{day}}$ | Daytime transpiration (ml) |
| $T_{\text{night}}$ | Nighttime transpiration (ml) |
| $b_3$ | Fractionation by Rubisco (30 ‰) |
| $b_4$ | Fractionation of PEP carboxylation and isotopic equilibrium during dissolution and hydration of $\text{CO}_2$ (−5.2 ‰ at a leaf temperature of 30 °C) |
| $S$ | Fractionation during the $\text{CO}_2$ leakage from the bundle sheath cells (1.8 ‰) |
| $\Phi$ | Leakiness of $\text{CO}_2$ from the bundle sheath |
| $H^2$ | Broad sense heritability |

