## Supplementary material for "A genetic link between leaf carbon isotope composition and whole-plant water use efficiency in the C_4_ grass *Setaria*"

### Supplementary Tables and Figures

**Table S1** Analysis of leaf composition traits.

| Trait | Treatment block | | | | | | Treatment | | Genotype | | Treatment x Genotype | |
|---|---|---|---|---|---|---|---|---|---|---|---|---|
| | Well-watered | | | Water-limited | | | | | | | | |
| | Mean ± SD | Min | Max | Mean ± SD | Min | Max | $F_{ddf,ndf}$ | $P$ | $F_{ddf,ndf}$ | $P$ | $F_{ddf,ndf}$ | $P$ |
| Leaf N (%) | 2.81 ± 0.84 | 1.29 | 5.2 | 3.73 ± 0.66 | 2.3 | 5.326 | $170.55_{1,60}$ | <0.001 | $1.84_{152,60}$ | 0.01 | $0.804_{140,60}$ | 0.85 |
| Leaf N (g N m$^{-2}$) | 1.12 ± 0.33 | 0.28 | 2.32 | 1.30 ± 0.32 | 0.21 | 2.22 | $72.57_{1,344}$ | <0.001 | $2.10_{184,344}$ | <0.001 | $1.146_{175,344}$ | 0.15 |
| Leaf C (g C m$^{-2}$) | 16.24 ± 4.55 | 3.54 | 32.34 | 14.92 ± 3.77 | 2.84 | 28.35 | $26.26_{1,60}$ | <0.001 | $2.685_{184,344}$ | <0.001 | $1.228_{175,344}$ | 0.06 |
| C:N ratio | 14.69 ± 5.01 | 4.88 | 30.97 | 12.47 ± 3.72 | 4.43 | 32.28 | $67.48_{1,351}$ | <0.001 | $2.267_{184,351}$ | <0.001 | $1.705_{175,351}$ | <0.001 |

Means ± SD were determined on day 34 when leaves were harvested.

**Table S2** Broad-sense heritability ($H^2$) and proportional variance of traits from leaves harvested on day 34.

| Trait | Proportional variance | | | $H^2$ | | |
|---|---|---|---|---|---|---|
| | Genotype | Treatment | G x Treatment | Both treatments | Well-watered treatment | Water-limited treatment |
| Leaf N (%) | 0.19 | 0.43 | 0.00 | 0.61 | 0.40 | 0.11 |
| Leaf N (g N/m$^{-2}$) | 0.16 | 0.13 | 0.06 | 0.41 | 0.38 | 0.08 |
| Leaf C (g C/m$^{-2}$) | 0.25 | 0.05 | 0.05 | 0.53 | 0.41 | 0.22 |
| C:N ratio | 0.15 | 0.07 | 0.17 | 0.36 | 0.43 | 0.27 |

**Table S3** A statistical comparison of additive and epistatic QTL models

| Trait | Tmt | Chr1 | Chr2 | pos1f | pos2f | lod.full | pval | lod.fv1 | pval.1 | lod.int | pval.2 | pos1a | pos2a | lod.add | pval.3 | lod.av1 | pval.4 |
|---|---|---|---|---|---|---|---|---|---|---|---|---|---|---|---|---|---|
| $T_{day}$ | WL | 4 | 7 | 56.40 | 99.94 | 7.00 | 0.05 | 3.83 | 0.51 | 0.03 | 1 | 52.90 | 99.94 | 6.97 | 0.01 | 3.80 | 0.03 |
| $T_{day}$ | WL | 4 | 9 | 56.40 | 33.94 | 11.77 | 0 | 3.75 | 0.56 | 0.07 | 1 | 56.40 | 33.94 | 11.70 | 0 | 3.67 | 0.03 |
| $T_{day}$ | WL | 5 | 9 | 115.86 | 33.94 | 13.21 | 0 | 5.19 | 0.04 | 0.07 | 1 | 115.86 | 33.94 | 13.14 | 0 | 5.12 | 0 |
| $T_{night}$ | WW | 6 | 7 | 87.12 | 65.07 | 8.07 | 0.01 | 3.65 | 0.66 | 0.30 | 1 | 90.75 | 65.07 | 7.77 | 0 | 3.35 | 0.04 |
| $T_{night}$ | WW | 7 | 9 | 65.07 | 34.93 | 8.50 | 0.01 | 4.07 | 0.39 | 0.00 | 1 | 65.07 | 34.93 | 8.49 | 0 | 4.07 | 0 |
| $T_{day}$ | WW | 2 | 5 | 94.71 | 109.70 | 7.61 | 0 | 3.32 | 0.85 | 0.08 | 1 | 98.51 | 104.13 | 7.52 | 0 | 3.24 | 0.02 |
| $T_{day}$ | WW | 2 | 7 | 98.51 | 99.94 | 11.15 | 0 | 5.66 | 0.03 | 0.41 | 1 | 98.51 | 99.94 | 10.74 | 0 | 5.25 | 0 |
| $T_{day}$ | WW | 2 | 9 | 98.51 | 34.93 | 14.13 | 0 | 6.06 | 0.01 | 0.37 | 1 | 98.51 | 34.93 | 13.75 | 0 | 5.68 | 0 |
| $T_{day}$ | WW | 5 | 7 | 109.70 | 99.94 | 10.03 | 0 | 4.54 | 0.13 | 1.30 | 1 | 80.93 | 99.94 | 8.73 | 0 | 3.24 | 0.02 |
| $T_{day}$ | WW | 5 | 9 | 80.93 | 34.93 | 12.75 | 0 | 4.68 | 0.11 | 0.38 | 1 | 80.93 | 34.93 | 12.36 | 0 | 4.29 | 0 |
| $T_{day}$ | WW | 7 | 9 | 99.94 | 33.94 | 14.22 | 0 | 6.15 | 0.01 | 0.08 | 1 | 99.94 | 33.94 | 14.14 | 0 | 6.07 | 0 |
| $T_{total}$ | WL | 2 | 7 | 108.89 | 94.06 | 8.45 | 0 | 3.77 | 0.4 | 0.88 | 1 | 98.51 | 99.94 | 7.57 | 0 | 2.89 | 0.05 |
| $T_{total}$ | WL | 2 | 9 | 98.51 | 34.93 | 13.90 | 0 | 3.69 | 0.46 | 0.01 | 1 | 98.51 | 34.93 | 13.89 | 0 | 3.68 | 0 |
| $T_{total}$ | WL | 4 | 7 | 52.90 | 99.94 | 7.87 | 0 | 3.18 | 0.85 | 0.06 | 1 | 52.90 | 99.94 | 7.81 | 0 | 3.13 | 0.02 |
| $T_{total}$ | WL | 5 | 9 | 115.13 | 34.93 | 13.82 | 0 | 3.60 | 0.52 | 0.26 | 1 | 115.86 | 34.93 | 13.55 | 0 | 3.34 | 0 |
| $T_{total}$ | WL | 7 | 9 | 96.48 | 34.93 | 16.74 | 0 | 6.53 | 0 | 0.24 | 1 | 96.48 | 34.93 | 16.50 | 0 | 6.28 | 0 |
| Biomass | WL | 2 | 9 | 98.51 | 33.94 | 13.53 | 0 | 6.22 | 0 | 0.01 | 1 | 98.51 | 33.94 | 13.52 | 0 | 6.21 | 0 |
| Biomass | WL | 5 | 9 | 80.93 | 33.94 | 11.20 | 0 | 3.89 | 0.58 | 0.19 | 1 | 115.86 | 33.94 | 11.00 | 0 | 3.70 | 0.05 |
| Biomass | WL | 9 | 9 | 33.94 | 126.28 | 11.29 | 0 | 3.98 | 0.5 | 0.26 | 1 | 33.94 | 126.28 | 11.03 | 0 | 3.73 | 0.05 |
| $T_{total}$ | WW | 2 | 5 | 96.05 | 109.70 | 8.34 | 0.01 | 3.77 | 0.52 | 0.03 | 1 | 96.05 | 109.70 | 8.31 | 0 | 3.74 | 0 |
| $T_{total}$ | WW | 2 | 7 | 97.69 | 99.94 | 11.38 | 0 | 5.80 | 0.01 | 0.00 | 1 | 97.69 | 99.94 | 11.38 | 0 | 5.80 | 0 |
| $T_{total}$ | WW | 2 | 9 | 95.91 | 34.93 | 14.49 | 0 | 7.00 | 0 | 0.67 | 1 | 95.91 | 34.93 | 13.82 | 0 | 6.33 | 0 |
| $T_{total}$ | WW | 5 | 9 | 80.93 | 34.93 | 10.82 | 0 | 3.33 | 0.81 | 0.39 | 1 | 80.93 | 34.93 | 10.43 | 0 | 2.94 | 0.05 |
| $T_{total}$ | WW | 7 | 9 | 99.94 | 34.93 | 13.76 | 0 | 6.28 | 0.01 | 0.01 | 1 | 99.94 | 34.93 | 13.76 | 0 | 6.27 | 0 |
| Biomass | WW | 2 | 5 | 95.91 | 89.13 | 11.25 | 0 | 4.88 | 0.03 | 0.47 | 1 | 95.91 | 80.93 | 10.78 | 0 | 4.41 | 0.01 |
| Biomass | WW | 2 | 7 | 97.69 | 99.94 | 11.40 | 0 | 5.03 | 0.01 | 0.09 | 1 | 97.69 | 99.94 | 11.31 | 0 | 4.94 | 0 |
| Biomass | WW | 2 | 9 | 98.51 | 34.93 | 14.22 | 0 | 7.85 | 0 | 0.04 | 1 | 98.51 | 34.93 | 14.17 | 0 | 7.80 | 0 |
| Biomass | WW | 3 | 5 | 52.18 | 79.49 | 7.98 | 0 | 3.36 | 0.75 | 0.05 | 1 | 52.18 | 80.93 | 7.93 | 0 | 3.31 | 0.04 |

| | | | | | | | | | | | | | | | | | |
|---|---|---|---|---|---|---|---|---|---|---|---|---|---|---|---|---|---|
| Biomass | WW | 3 | 7 | 52.18 | 58.88 | 7.23 | 0.01 | 3.52 | 0.62 | 0.09 | 1 | 52.18 | 99.94 | 7.14 | 0.01 | 3.43 | 0.04 |
| Biomass | WW | 5 | 6 | 80.93 | 80.46 | 7.62 | 0.01 | 2.99 | 0.97 | 0.00 | 1 | 80.93 | 80.46 | 7.62 | 0.01 | 2.99 | 0.04 |
| Biomass | WW | 5 | 7 | 80.93 | 99.94 | 8.67 | 0 | 4.05 | 0.3 | 0.60 | 1 | 80.93 | 99.94 | 8.07 | 0 | 3.45 | 0.04 |
| Biomass | WW | 5 | 9 | 80.93 | 34.93 | 12.44 | 0 | 6.47 | 0 | 0.82 | 1 | 80.93 | 34.93 | 11.63 | 0 | 5.65 | 0 |
| Biomass | WW | 7 | 9 | 96.02 | 34.93 | 10.28 | 0 | 4.30 | 0.19 | 0.04 | 1 | 96.02 | 34.93 | 10.23 | 0 | 4.26 | 0.01 |
| $WUE_{plant}$ | WW | 9 | 9 | 31.07 | 31.99 | 18.85 | 0 | 13.32 | 0.1 | 0.34 | 1 | 31.07 | 31.99 | 18.51 | 0 | 12.97 | 0 |
| $WUE_{plant}$ | WL | 7 | 9 | 94.06 | 34.93 | 9.55 | 0 | 4.27 | 0.24 | 0.01 | 1 | 94.06 | 34.93 | 9.54 | 0 | 4.26 | 0.01 |
| $WUE_{plant}$ | WW | 1 | 7 | 4.94 | 99.94 | 11.06 | 0 | 3.62 | 0.6 | 0.04 | 1 | 4.94 | 99.94 | 11.02 | 0 | 3.58 | 0.02 |
| $WUE_{plant}$ | WW | 7 | 9 | 99.94 | 34.93 | 15.60 | 0 | 8.16 | 0 | 0.03 | 1 | 99.94 | 34.93 | 15.57 | 0 | 8.13 | 0 |
| $\delta^{13}C_{leaf}$ | WW | 4 | 7 | 57.17 | 99.94 | 8.20 | 0.01 | 3.41 | 0.71 | 0.24 | 1 | 57.17 | 99.94 | 7.96 | 0 | 3.17 | 0.01 |
| $\delta^{13}C_{leaf}$ | WW | 7 | 7 | 51.65 | 99.94 | 8.57 | 0.01 | 3.78 | 0.51 | 0.09 | 1 | 51.65 | 99.94 | 8.48 | 0 | 3.69 | 0 |
| $\delta^{13}C_{leaf}$ | WW | 7 | 9 | 99.94 | 34.93 | 12.94 | 0 | 6.16 | 0 | 0.88 | 1 | 99.94 | 34.93 | 12.06 | 0 | 5.27 | 0 |

The statistical support of five QTL model components is presented. These models include ones that: (1) fit the additive and epistatic effects (lod.full), a comparison of the full model to one that includes additive effects of a single QTL with the possibility of a second locus that contributes epistatic effects (lod.fv1), (2) fit additive effects of a single major QTL relative to a model with additional QTL that exhibit no epistasis (lod.av1), (3) fit for a simple additive QTL model (lod.add), and (4) a comparison between the full model and the additive model (lod.int). The position of each QTL is denoted by both the chromosome (chr) and position (pos) where the position denoted by the full model and the additive model contain a "f' (pos.f) or "a" (pos.a), respectively. The statistical significance of each component is denoted by p-value directly to the left of each LOD score column (full.lod). Metadata describing trait, treatment are also included in the table. $T_{day}$, $T_{night}$, $T_{total}$ are daytime, nighttime, and total transpiration, respectively, and the treatment (tmt) are well-watered (WW) and water-limited (WL).

**Table S4** QTL found in all traits in both treatments.

| QTL | Chr. | Position | Trait | Treatment | LOD | Proportional Variance | Confidence Interval |
|---|---|---|---|---|---|---|---|
| 1@15 | 1 | 12.92 | WUEplant | Well-watered | 3.7 | 7.3 | 0 - 18.27 |
| 1@75 | 1 | 74.97 | WUEplant | Water-limited | 4.8 | -9.6 | 70.73 - 85.17 |
| 2@11 | 2 | 11.41 | Biomass | Well-watered | 4.5 | -5.4 | 8.4 - 16.33 |
| 2@92 | 2 | 97.45 | Biomass | Water-limited | 7.4 | 11.4 | 93.52 - 105.63 |
| 2@92 | 2 | 96.83 | Biomass | Well-watered | 10.7 | 16.6 | 94.78 - 100.27 |
| 2@92 | 2 | 96.30 | C:N ratio | Well-watered | 4.2 | 8.4 | 84.61 - 113.85 |
| 2@92 | 2 | 98.51 | Day transpiration | Well-watered | 7.1 | 10.7 | 95.22 - 108.89 |
| 2@92 | 2 | 96.30 | Leaf C | Well-watered | 4.5 | 8.9 | 84.61 - 98.51 |
| 2@92 | 2 | 90.65 | Leaf N (%) | Well-watered | 7.1 | -11.2 | 87.02 - 92.82 |
| 2@92 | 2 | 98.48 | Total transpiration | Water-limited | 5.2 | 8.7 | 93.74 - 106.71 |
| 2@92 | 2 | 96.73 | Total transpiration | Well-watered | 9.0 | 13.3 | 93.13 - 99.08 |
| 2@113 | 2 | 106.34 | Biomass | Well-watered | 5.4 | 11.0 | 93.53 - 113.85 |
| 2@113 | 2 | 106.34 | Day transpiration | Well-watered | 6.5 | 8.8 | 98.1 - 111.54 |
| 3@16 | 3 | 16.46 | Biomass | Well-watered | 2.9 | -3.5 | 0 - 86.56 |
| 3@49 | 3 | 47.98 | Biomass | Well-watered | 4.2 | 5.4 | 45.17 - 56.02 |
| 3@49 | 3 | 47.05 | Total transpiration | Well-watered | 5.0 | 5.7 | 44.12 - 47.39 |
| 3@64 | 3 | 64.48 | WUEplant | Water-limited | 3.9 | 7.6 | 61.06 - 84.64 |
| 3@84 | 3 | 84.64 | WUEplant | Water-limited | 3.2 | 5.9 | 69.68 - 104.08 |
| 4@50 | 4 | 50.73 | Biomass | Water-limited | 4.7 | 11.7 | 47.81 - 60.63 |
| 4@50 | 4 | 56.40 | Day transpiration | Water-limited | 3.8 | 5.7 | 47.81 - 59.59 |
| 4@50 | 4 | 52.90 | Total transpiration | Water-limited | 4.6 | 8.6 | 45.87 - 61 |
| 5@79 | 5 | 79.96 | Biomass | Water-limited | 4.8 | -8.0 | 71.92 - 84.31 |
| 5@79 | 5 | 86.50 | Biomass | Well-watered | 6.6 | -9.4 | 79.93 - 96.55 |
| 5@79 | 5 | 82.92 | Day transpiration | Well-watered | 4.7 | -6.5 | 74.1 - 99.77 |
| 5@79 | 5 | 78.65 | Leaf N (%) | Well-watered | 5.3 | 8.1 | 71.03 - 87.89 |

| | | | | | | | |
|---|---|---|---|---|---|---|---|
| 5@79 | 5 | 79.96 | WUEplant | Well-watered | 3.5 | -8.8 | 64.47 - 118.9 |
| 5@104 | 5 | 107.16 | Biomass | Water-limited | 5.6 | -8.3 | 92.89 - 119 |
| 5@104 | 5 | 105.80 | Biomass | Well-watered | 8.2 | -11.2 | 99.87 - 111.91 |
| 5@104 | 5 | 115.86 | Day transpiration | Water-limited | 5.4 | -9.8 | 106.54 - 120 |
| 5@104 | 5 | 115.13 | Day transpiration | Well-watered | 4.4 | -5.0 | 112.94 - 120.98 |
| 5@104 | 5 | 100.66 | Leaf C | Well-watered | 3.8 | -7.5 | 95.23 - 104.13 |
| 5@104 | 5 | 100.41 | Leaf N | Well-watered | 3.4 | -7.9 | 95.23 - 104.13 |
| 5@104 | 5 | 115.13 | Total transpiration | Water-limited | 4.4 | -7.0 | 110.84 - 120.24 |
| 5@104 | 5 | 109.33 | Total transpiration | Well-watered | 6.6 | -8.7 | 105.09 - 113.63 |
| 5@104 | 5 | 92.71 | WUEplant | Water-limited | 4.0 | -10.5 | 77.28 - 97.52 |
| 6@59 | 6 | 59.89 | Biomass | Well-watered | 4.3 | 4.7 | 56.47 - 65.46 |
| 6@59 | 6 | 61.39 | Day transpiration | Water-limited | 3.1 | 4.6 | 30.64 - 81.7 |
| 6@59 | 6 | 60.39 | Total transpiration | Well-watered | 3.7 | 4.2 | 51.1 - 78.52 |
| 6@59 | 6 | 49.08 | WUEplant | Well-watered | 3.5 | 8.2 | 44.73 - 58.82 |
| 6@75 | 6 | 73.67 | Biomass | Well-watered | 4.8 | 5.6 | 58.28 - 79.34 |
| 6@75 | 6 | 72.84 | Day transpiration | Well-watered | 3.9 | 4.4 | 62.38 - 80.21 |
| 6@75 | 6 | 72.84 | Leaf N (%) | Well-watered | 3.7 | -5.6 | 68.73 - 92.02 |
| 6@75 | 6 | 85.47 | Night transpiration | Well-watered | 3.2 | 5.6 | 80.21 - 92.02 |
| 7@33 | 7 | 33.82 | Biomass | Well-watered | 3.7 | -4.5 | 7.83 - 45.37 |
| 7@33 | 7 | 33.55 | Total transpiration | Well-watered | 3.7 | -4.7 | 17.74 - 54.91 |
| 7@33 | 7 | 25.14 | WUEplant | Well-watered | 3.5 | 5.7 | 19.91 - 81.23 |
| 7@51 | 7 | 50.95 | Biomass | Water-limited | 5.2 | -7.3 | 50.24 - 51.41 |
| 7@51 | 7 | 50.95 | Biomass | Well-watered | 4.8 | -6.9 | 42.81 - 58.1 |
| 7@51 | 7 | 51.18 | d13C | Well-watered | 3.5 | -6.5 | 40.98 - 70.05 |
| 7@51 | 7 | 51.18 | Day transpiration | Well-watered | 5.9 | -6.9 | 46.24 - 51.87 |
| 7@51 | 7 | 64.29 | Leaf N | Well-watered | 3.6 | -8.5 | 51.65 - 101.95 |
| 7@51 | 7 | 58.88 | Leaf N (%) | Well-watered | 4.8 | 7.3 | 54.73 - 70.05 |
| 7@51 | 7 | 65.02 | Night transpiration | Well-watered | 5.0 | -10.5 | 60.87 - 72.08 |

| | | | | | | | |
|---|---|---|---|---|---|---|---|
| 7@51 | 7 | 51.18 | Total transpiration | Well-watered | 5.0 | -5.9 | 36.14 - 54.57 |
| 7@77 | 7 | 77.36 | WUEplant | Well-watered | 3.5 | 7.5 | 61.15 - 92.51 |
| 7@99 | 7 | 96.75 | Biomass | Water-limited | 4.6 | -6.8 | 88.15 - 101.78 |
| 7@99 | 7 | 99.11 | Biomass | Well-watered | 6.6 | -9.4 | 95.16 - 101.95 |
| 7@99 | 7 | 99.94 | C:N ratio | Well-watered | 5.1 | -10.4 | 96.48 - 101.95 |
| 7@99 | 7 | 99.94 | d13C | Well-watered | 4.3 | -8.2 | 96.48 - 101.95 |
| 7@99 | 7 | 96.02 | Day transpiration | Water-limited | 3.7 | -6.2 | 89.93 - 101.95 |
| 7@99 | 7 | 97.14 | Day transpiration | Well-watered | 6.3 | -9.0 | 91.78 - 101.38 |
| 7@99 | 7 | 99.94 | Leaf C | Well-watered | 4.8 | -9.6 | 62.63 - 101.95 |
| 7@99 | 7 | 99.94 | Leaf N (%) | Well-watered | 7.3 | 11.6 | 96.48 - 101.95 |
| 7@99 | 7 | 98.17 | Total transpiration | Water-limited | 6.5 | -11.2 | 90.97 - 101.48 |
| 7@99 | 7 | 99.94 | Total transpiration | Well-watered | 6.8 | -10.0 | 88.8 - 102.14 |
| 7@99 | 7 | 94.65 | WUEplant | Water-limited | 4.1 | 8.6 | 90 - 101.55 |
| 7@99 | 7 | 96.66 | WUEplant | Well-watered | 7.5 | 15.1 | 89.66 - 102.32 |
| 9@33 | 9 | 33.49 | Biomass | Water-limited | 11.4 | 18.3 | 31.85 - 37.61 |
| 9@33 | 9 | 34.78 | Biomass | Well-watered | 6.3 | 9.2 | 31.86 - 38.75 |
| 9@33 | 9 | 33.94 | C:N ratio | Well-watered | 5.8 | 11.8 | 31.53 - 36.41 |
| 9@33 | 9 | 34.93 | d13C | Well-watered | 7.4 | 14.5 | 30.61 - 38.06 |
| 9@33 | 9 | 33.94 | Day transpiration | Water-limited | 10.7 | 20.7 | 31.99 - 35.85 |
| 9@33 | 9 | 33.46 | Day transpiration | Well-watered | 10.6 | 15.9 | 29.95 - 34.67 |
| 9@33 | 9 | 22.74 | Leaf C | Well-watered | 3.3 | 6.5 | 19.9 - 83.97 |
| 9@33 | 9 | 34.93 | Leaf N (%) | Well-watered | 7.1 | -11.3 | 31.53 - 37.12 |
| 9@33 | 9 | 35.91 | Night transpiration | Water-limited | 3.8 | 9.7 | 32.79 - 38.06 |
| 9@33 | 9 | 33.85 | Night transpiration | Well-watered | 4.8 | 10.9 | 30.75 - 35.95 |
| 9@33 | 9 | 35.10 | Total transpiration | Water-limited | 12.2 | 22.5 | 32.71 - 37.41 |
| 9@33 | 9 | 35.08 | Total transpiration | Well-watered | 10.8 | 16.1 | 33.07 - 37.53 |
| 9@33 | 9 | 36.59 | WUEplant | Water-limited | 7.2 | -16.1 | 32.29 - 38.77 |
| 9@33 | 9 | 34.98 | WUEplant | Well-watered | 8.7 | -17.1 | 32.79 - 37.32 |

| | | | | | | | |
|---|---|---|---|---|---|---|---|
| 9@33 | 9 | 93.48 | Night transpiration | Well-watered | 3.2 | -5.6 | 47.69 - 165.58 |
| 9@126 | 9 | 126.11 | Biomass | Water-limited | 5.0 | 7.7 | 110.08 - 133.25 |
| 9@126 | 9 | 132.18 | Biomass | Well-watered | 4.3 | 6.8 | 126.5 - 141.1 |

Position, LOD, Variance, and confidence intervals are means. Confidence intervals are for the mean position of the QTL. Proportional variance is the proportion of additive variance explained (%) by the QTL, which can have a positive or negative effect on the trait.

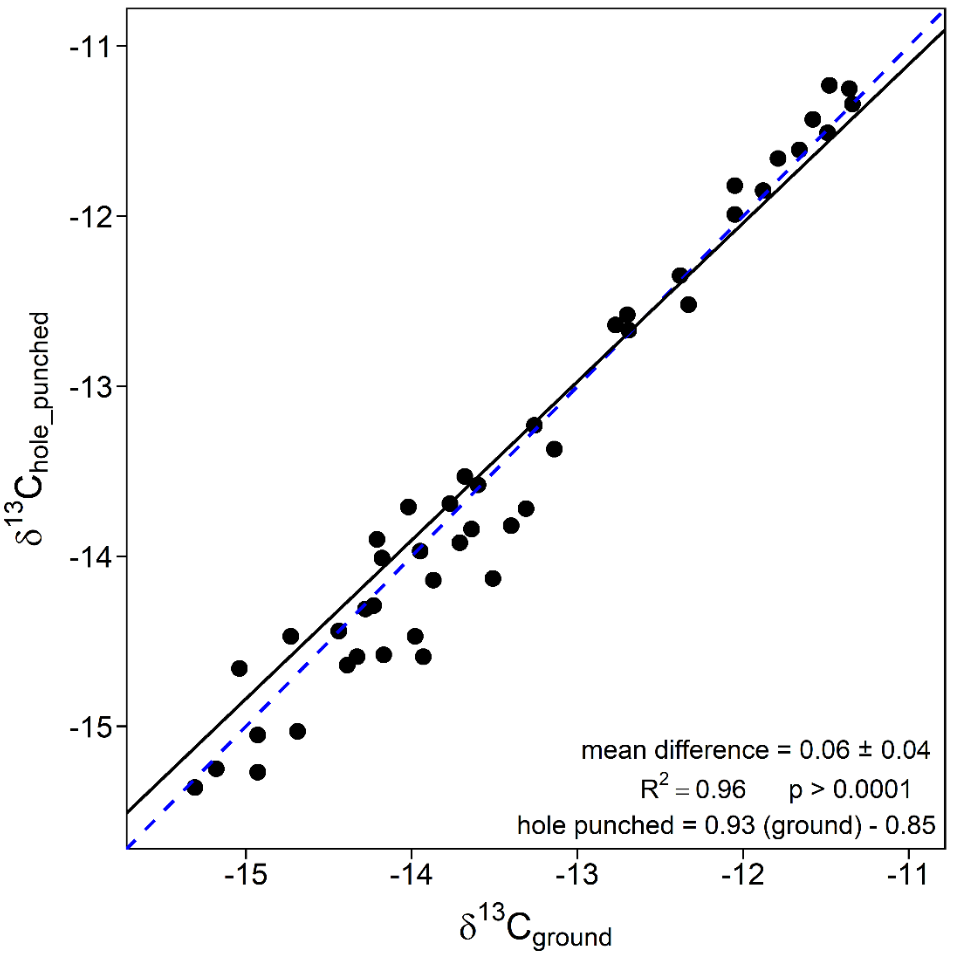


**Fig. S1** Comparison of the two sampling methods for $\delta^{13}C_{leaf}$. Principal method used to sample for $\delta^{13}C_{leaf}$ was hole punching, which provided similar values to grinding the entire leaf and weighing out a subsample. Solid, black line represents the regression line. The dashed, blue represents 1:1 line.

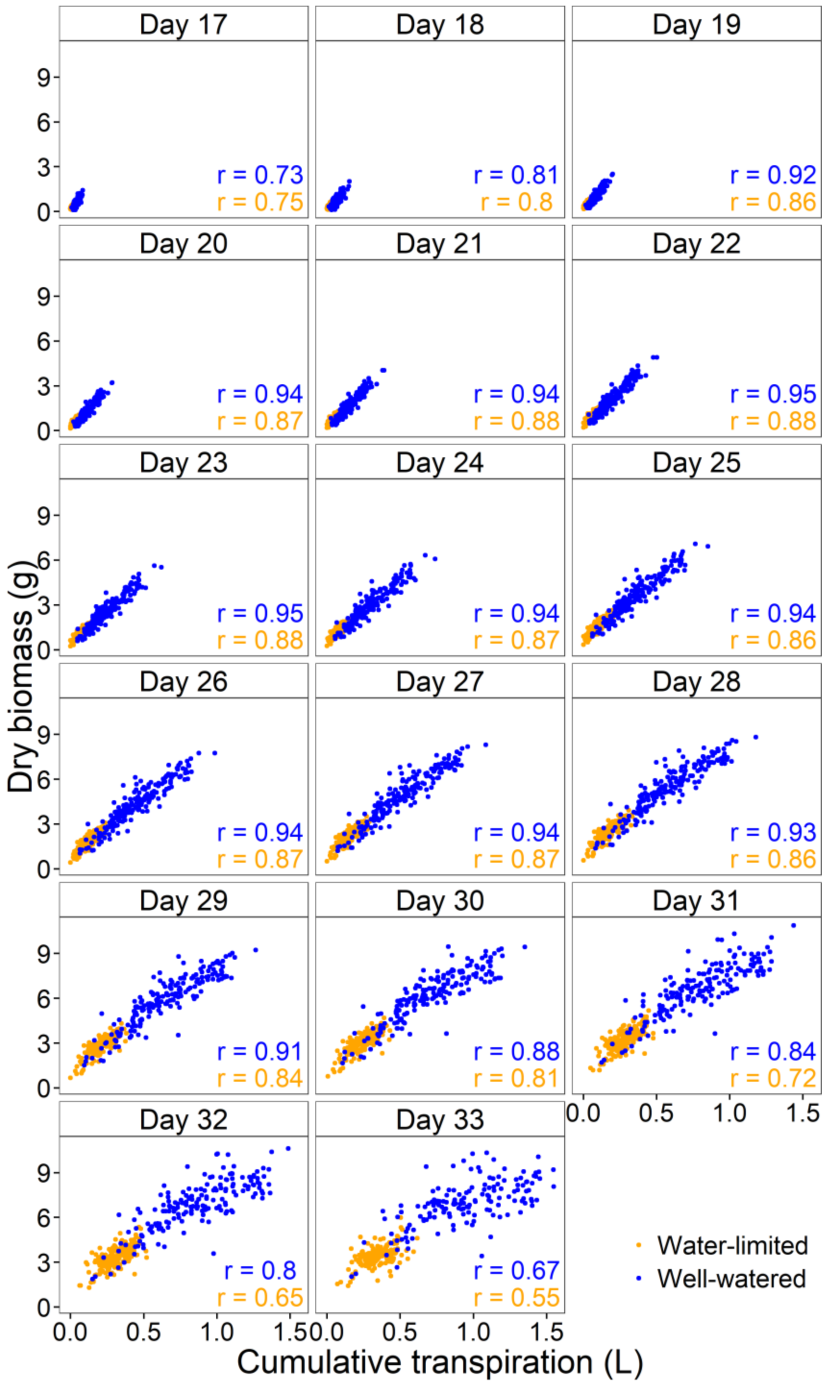


**Fig. S2** Dry biomass versus cumulative transpiration for each day of the experiment. Correlations for each treatment are given in each figure.

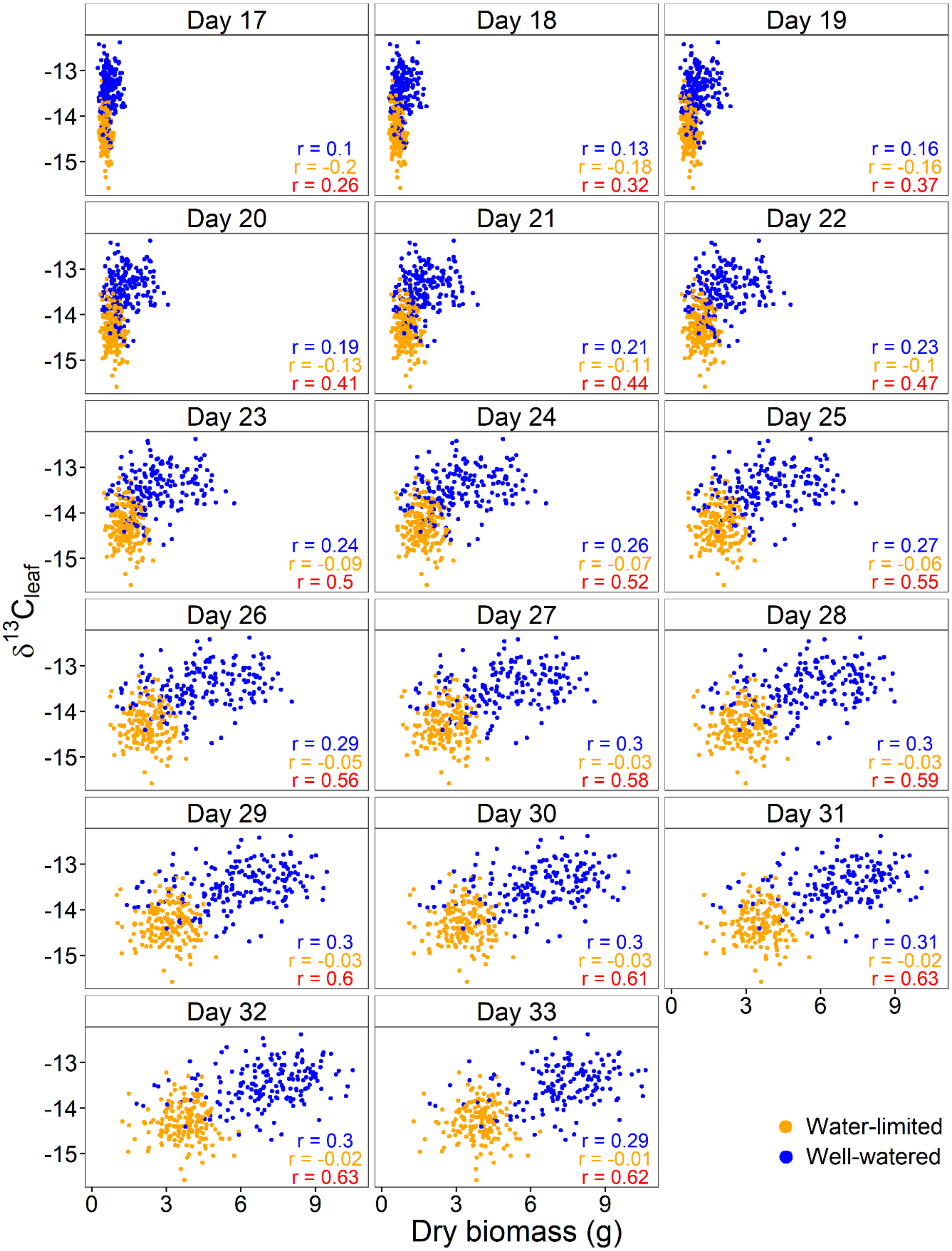


**Fig. S3** Relationship between $\delta^{13}C_{leaf}$ and dry aboveground biomass for each day of the experiment. Correlations are given in each panel.

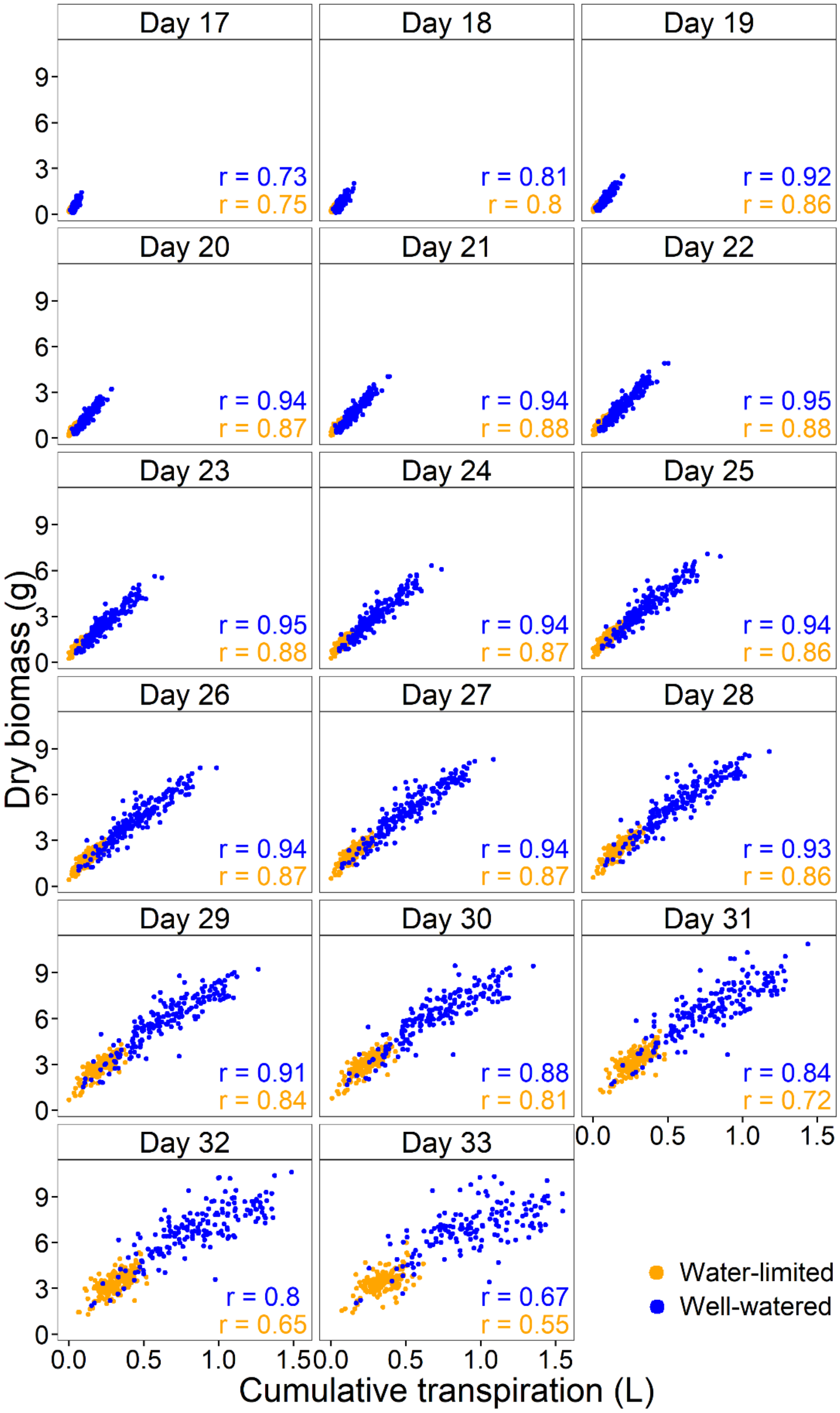


**Fig. S4** Relationship between $\delta^{13}C_{leaf}$ and cumulative transpiration for each day of the experiment. Correlations are given in each panel.

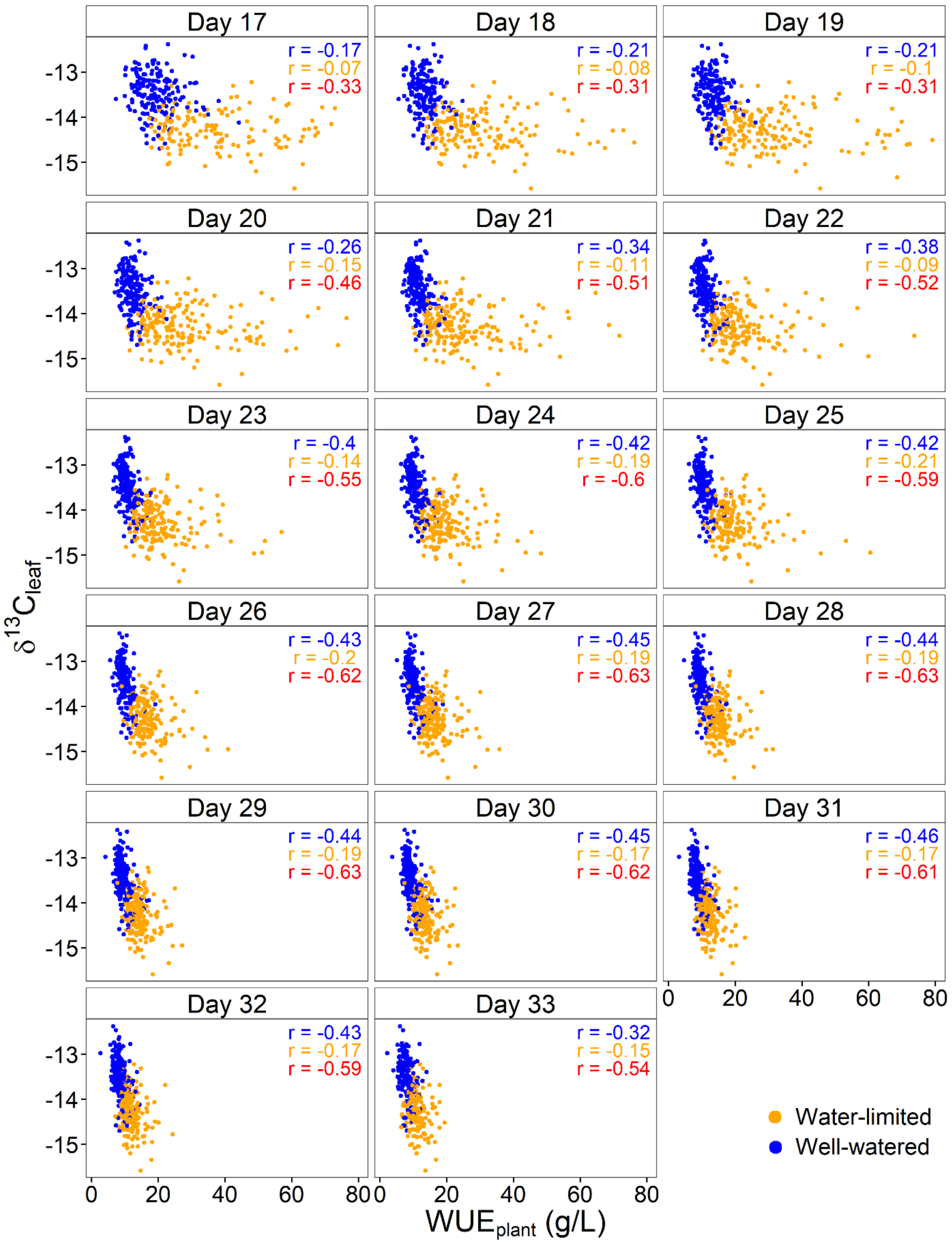


**Fig. S5** Relationship between $\delta^{13}C_{leaf}$ and $WUE_{plant}$ for each day of the experiment. Correlations in each treatment are given in each panel.

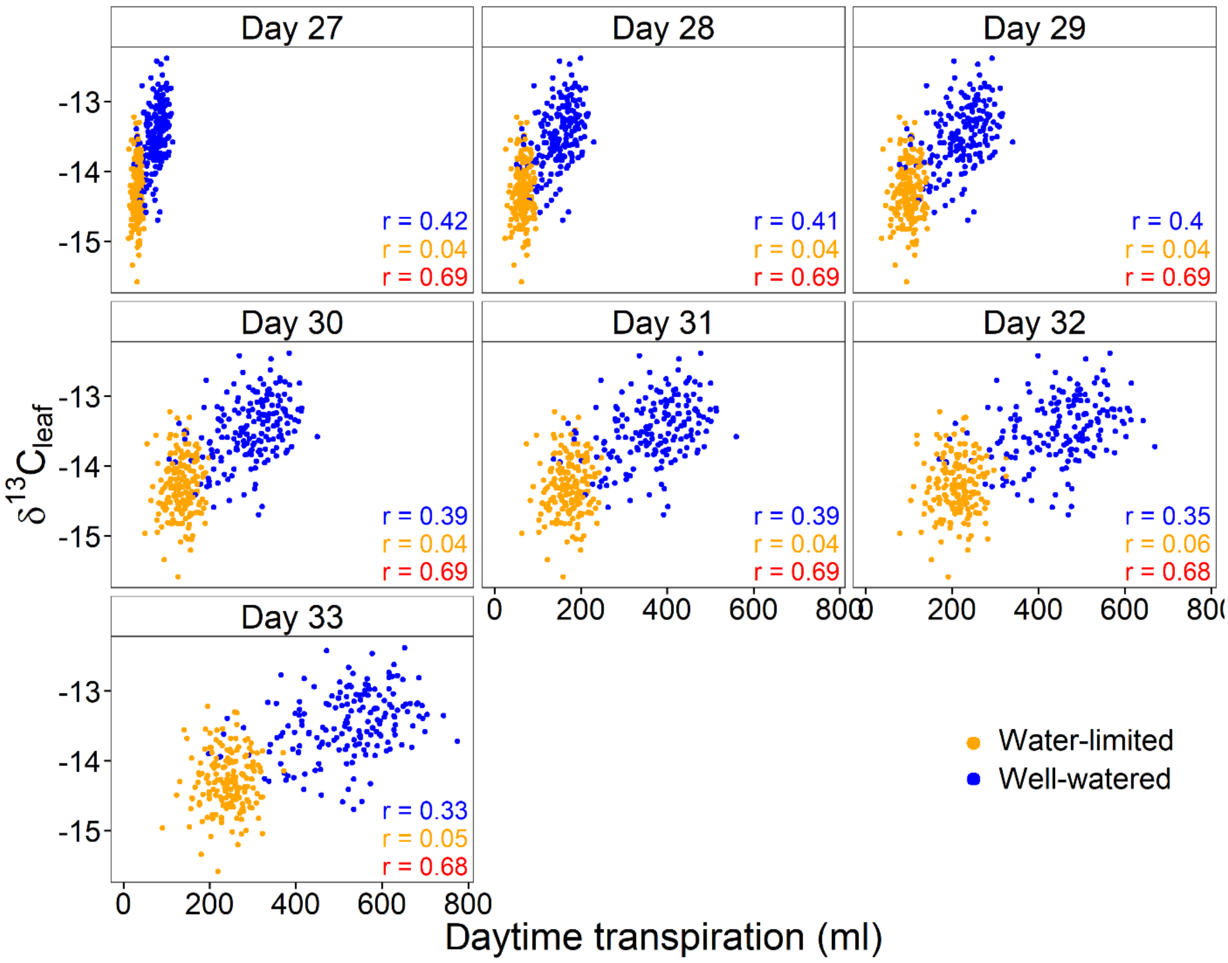


**Fig. S6** Relationship between $\delta^{13}C_{leaf}$ and $T_{day}$ for each day of the experiment. Correlations in each treatment are given in each panel.

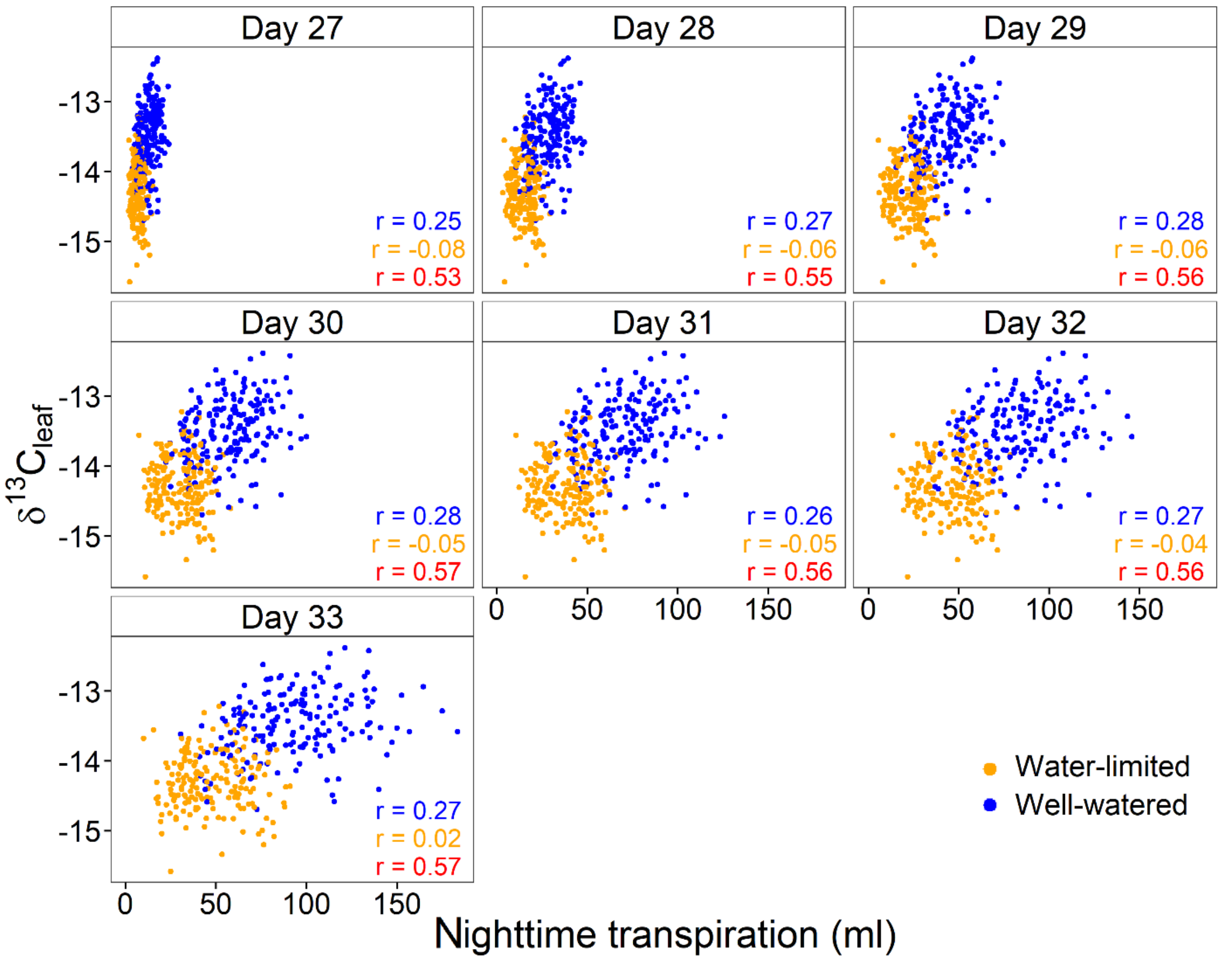


**Fig. S7** Relationship between $\delta^{13}C_{leaf}$ and $T_{night}$ for each day of the experiment. Correlations in each treatment are given in each panel.

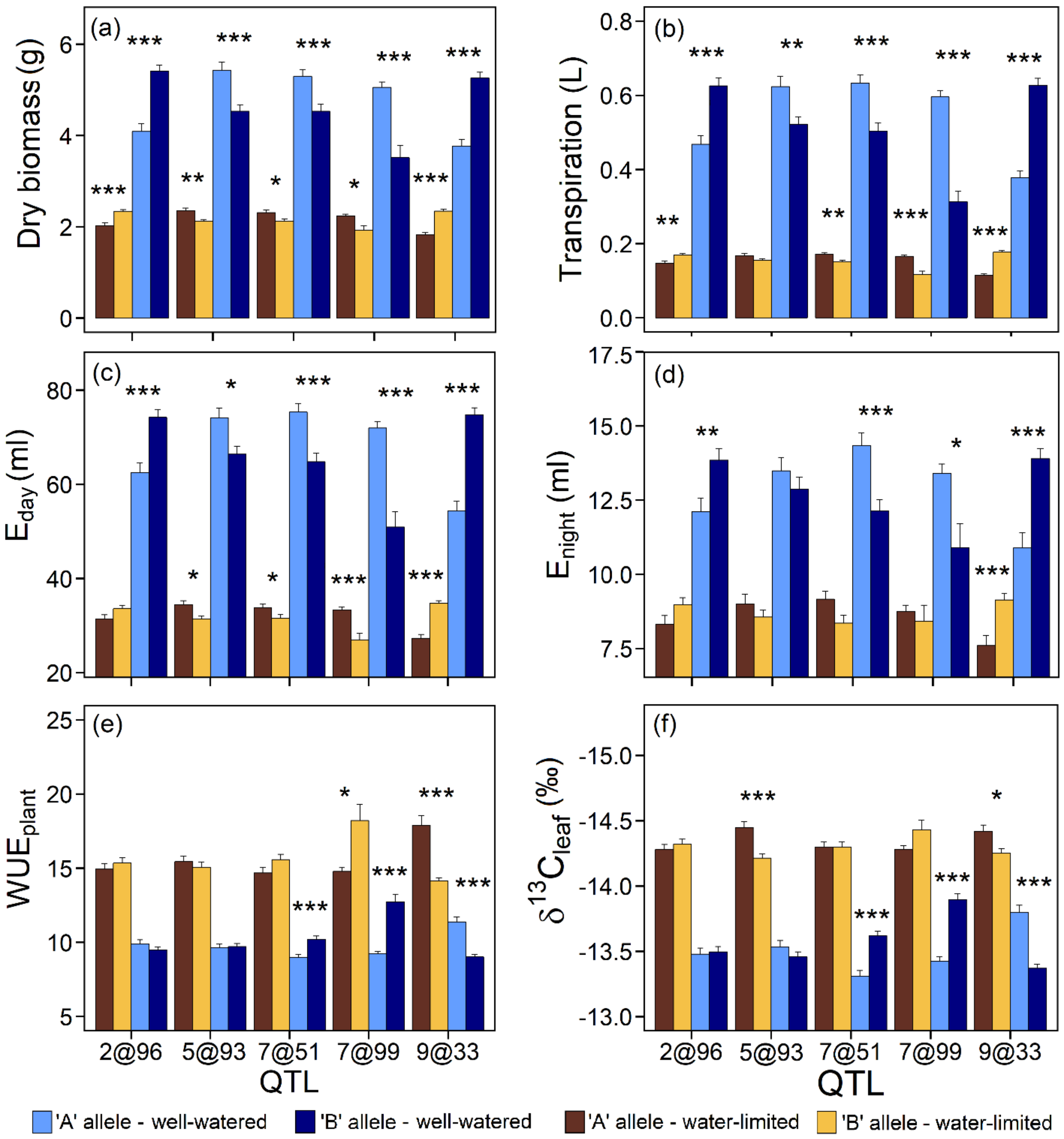


**Fig. S8** Effect of alleles of the five principal QTL identified for dry biomass (a), transpiration (b), $T_{day}$ (c), $T_{night}$ (d), $WUE_{plant}$ (e), $\delta^{13}C_{leaf}$ (f). 'A' represents the allele from the A10 parental line (*Setaria viridis*), and 'B' represents the allele from the B100 parental line (*Setaria italica*). Level of significance is denoted as the following: *, **, *** represent P < 0.05, 0.01, 0.0001, respectively
